# Differential metabolism of arsenicals regulates Fps1-mediated arsenite transport

**DOI:** 10.1101/2021.10.12.464049

**Authors:** Jongmin Lee, David E. Levin

## Abstract

Arsenic is an environmental toxin that exists mainly as pentavalent arsenate and trivalent arsenite. Both forms activate the yeast SAPK Hog1, but with different consequences. We describe a mechanism by which cells distinguish between these arsenicals through one-step metabolism to differentially regulate the bidirectional glycerol channel Fps1, an adventitious port for arsenite. Cells exposed to arsenate reduce it to thiol-reactive arsenite, which modifies a set of cysteine residues in target proteins; whereas cells exposed to arsenite metabolize it to methylarsenite, which modifies an additional set of cysteine residues. Hog1 becomes arsenylated, which prevents it from closing Fps1. However, this block is overcome in cells exposed to arsenite through methylarsenylation of Acr3, an arsenite efflux pump that we found also regulates Fps1 directly. This adaptation allows cells to restrict arsenite entry through Fps1, but also allows its exit when produced from arsenate exposure. These results have broad implications for understanding how SAPKs activated by diverse stressors can drive stress-specific outputs.

**Summary** (for JCB table of contents)

Jongmin Lee, David E. Levin

Lee and Levin investigate the question of how different stressors can drive divergent outputs from an active SAPK. The work describes the mechanism by which two forms of arsenic that both activate the yeast SAPK Hog1 differentially regulate the glycerol channel Fps1.

## Introduction

Arsenic is the most prevalent toxin in the environment (Rosen and Liu, 2009). This natural metalloid enters the biosphere from geochemical sources and, to a lesser degree, from anthropogenic sources (Zhu et al., 2014). Its ubiquitous presence has driven the evolution of arsenic resistance mechanisms, which exist in nearly every organism (Yang and Rosen, 2016). Human exposure to arsenic is mainly through food, water and air, and contamination of groundwater is a worldwide health problem (Smedley and Kinniburgh, 2002; Naujokas et al., 2013). Environmental arsenic exists mainly as oxyanions of inorganic trivalent arsenite [As(III)] and pentavalent arsenate [As(V)]. As(V) is much less toxic than As(III), which is thiol-reactive and binds covalently to cysteine residues in proteins (Shen et al., 2013). Chronic exposure to inorganic arsenic is associated with cardiovascular disease and hypertension, diabetes mellitus, neurological disorders, and various forms of cancer (Abernathy et al., 2003; Beane Freeman et al., 2004; Naujokas et al., 2013). It has been proposed that both direct modification of biomolecules by As(III) and reactive oxygen species generated by arsenicals are responsible for their toxicity and carcinogenicity (Hughes et al., 2011; Martinez et al., 2011). Despite these health effects, arsenic trioxide [which solubilizes to As(III)] is used as a highly effective treatment for acute promyelocytic leukemia (Liu et al., 2012; Kozono et al., 2018). Thus, it is important to understand the cellular responses mobilized by arsenic exposure.

In mammalian cells, uptake of As(V), which is an analog of inorganic phosphate, is mediated by the high-affinity phosphate transporter NaPi-IIb (Maciaszczyk-Dziubinska et al., 2012). Once inside the cell, As(V) must be reduced to As(III) by isoforms B and C of Cdc25 protein phosphatases/As(V) reductases for its elimination (Bhattacharjee et al., 2010). As(III), on the other hand, enters cells through the aquaglyceroporins and the glucose permeases (Maciaszczyk-Dziubinska et al., 2012). It can be transported out of the cell directly, or after conjugation with glutathione, by various ABC family transporters. Additionally, As(III) is metabolized in mammals by the As(III) S-adenosylmethionine (SAM) methyltransferase (AS3MT), which catalyzes the transfer of methyl groups from SAM to As(III), to produce methylarsenite [MAs(III)] and dimethylarsenite [DMA(III)] (Cullen, 2014; Dheeman et al., 2014), both of which are more toxic than inorganic As(III) and are similarly thiol-reactive (Styblo et al., 2000; Dong et al., 2015). It is thought that conversion of As(III) to MAs(III) and DMA(III) in humans assists in its excretion in the urine (Vahter and Concha, 2001; Dong et al., 2015; Jansen et al., 2016). However, our recent work in yeast reveals that production of MAs(III) from As(III) is also a metabolic activation step required to mobilize a coherent cellular response to As(III) (Lee and Levin, 2018; 2019; Lee et al., 2019).

In the yeast *S. cerevisiae*, As(V) uptake is through the phosphate transporters (Persson et al., 1998; Wysocki and Tamas, 2010). As(V) is then reduced to As(III) by the As(V) reductase Acr2 (Mukhopadhyay and Rosen, 1998). Similarly to mammalian cells, As(III) enters yeast cells principally through the aquaglyceroporin Fps1, a bidirectional channel that normally functions to transport glycerol (Wysocki et al., 2001), and secondarily through the hexose permeases (Maciaszczyk-Dziubinska et al., 2012). As(III) is actively transported out of the yeast cell through the plasma membrane metalloid/H+ antiporter, Acr3 (Maciaszczyk-Dziubinska et al., 2011). Alternatively, As(III) can be conjugated to glutathione and sequestered in the yeast vacuole through ABC family transporters Ycf1 and Vmr1 (Maciaszczyk-Dziubinska et al., 2012). We demonstrated recently that As(III) is converted to MAs(III) by the dimeric methyltransferase Mtq2:Trm112 (Lee and Levin, 2018).

The transcriptional response to arsenic in yeast is carried out principally by the AP-1-like transcription factor, Yap8/Acr1, a highly specialized transcriptional regulator that induces the expression of just two genes, *ACR2* and *ACR3*, which are transcribed in opposite directions from a common promoter (Wysocki et al., 2004). Yap8 is an As(III) sensor whose modification of three cysteine residues by As(III) converts it to an active transcriptional regulator (Kumar et al., 2015).

With regard to signaling, As(III) stimulates the mammalian SAPK p38 (Elbirt et al., 1998; Verma et al., 2002). The yeast ortholog of p38, Hog1 (Han et al., 1994), is similarly activated in response to As(III) treatment and plays an important role in the tolerance to As(III) (Sotelo and Rodríguez-Gabriel, 2006; Thorsen et al., 2006).

Activation of Hog1 by As(III) occurs through an indirect route that involves its metabolic activation to MAs(III) (Lee and Levin, 2018). MAs(III) inhibits the tyrosine-specific protein phosphatases, Ptp2 and Ptp3, which normally maintain Hog1 in a low-activity state. Active Hog1 protects cells from As(III) toxicity, in part, by inducing closure of Fps1 to restrict its entry (Thorsen et al., 2006). This occurs when Hog1 phosphorylates the redundant regulators of the glycerol channel, Rgc1 and Rgc2, which drives their displacement from Fps1 similarly to hyper-osmotic stress (Lee et al., 2013, Lee and Levin, 2018). In the latter case, Hog1 closes Fps1 to restore osmotic balance through the accumulation of glycerol. However, glycerol accumulation is prevented in response to As(III) treatment, despite the closure of Fps1, through direct inhibition by MAs(III) of glycerol-3-phosphate dehydrogenase, the first committed step in glycerol biosynthesis from glycolytic intermediates (Lee and Levin, 2019). This adaptation allows the cell to avoid the osmotic imbalance that would otherwise result from inappropriate accumulation of glycerol while Fps1 is closed.

As(V) also activates Hog1, but through a different route that does not require its reduction to As(III) by Acr2, and involves activation of the MEK Pbs2 (Lee and Levin, 2018). However, activation of Hog1 in response to an As(V) challenge does not result in the closure of Fps1. This makes sense from a physiologic perspective, because As(V) enters the cell through the phosphate transporters rather than through Fps1, thus the bidirectional channel activity of Fps1 is important to export As(III) that is produced from the metabolism of As(V) (Maciaszczyk-Dziubinska et al., 2012). Because many microorganisms convert one arsenic oxidation state to the other (Muller et al., 2007; Wang et al., 2015), yeast in the wild may be exposed to either form in high concentration.

This study was motivated by our desire to understand the mechanisms by which various stress signals that activate a common SAPK elicit stress-specific outputs under the control of the activated protein kinase. In this case, we sought to understand how cells differentially regulate Fps1 in response to an As(III) challenge or an As(V) challenge. We describe an arsenic stress signaling code that explains how cells distinguish between these stressors to mount responses appropriate to each.

## Results

Although Hog1 is activated in cells exposed to either As(III) or As(V), important differences exist in the responses to these related stressors (Lee and Levin, 2018). For example, the cell closes the bidirectional glycerol channel Fps1 in response to As(III) challenge to restrict its entry to the cell, but leaves it open in response to As(V) challenge so as to allow exit of the As(III) produced. However, because As(V) must be reduced to As(III) before it is eliminated (Figure 1A; Maciaszczyk-Dziubinska et al., 2012; Garbinski et al., 2019), the cell is faced with the problem of distinguishing an As(III) challenge from an As(V) challenge with subsequent production of As(III).

**Figure 1.**
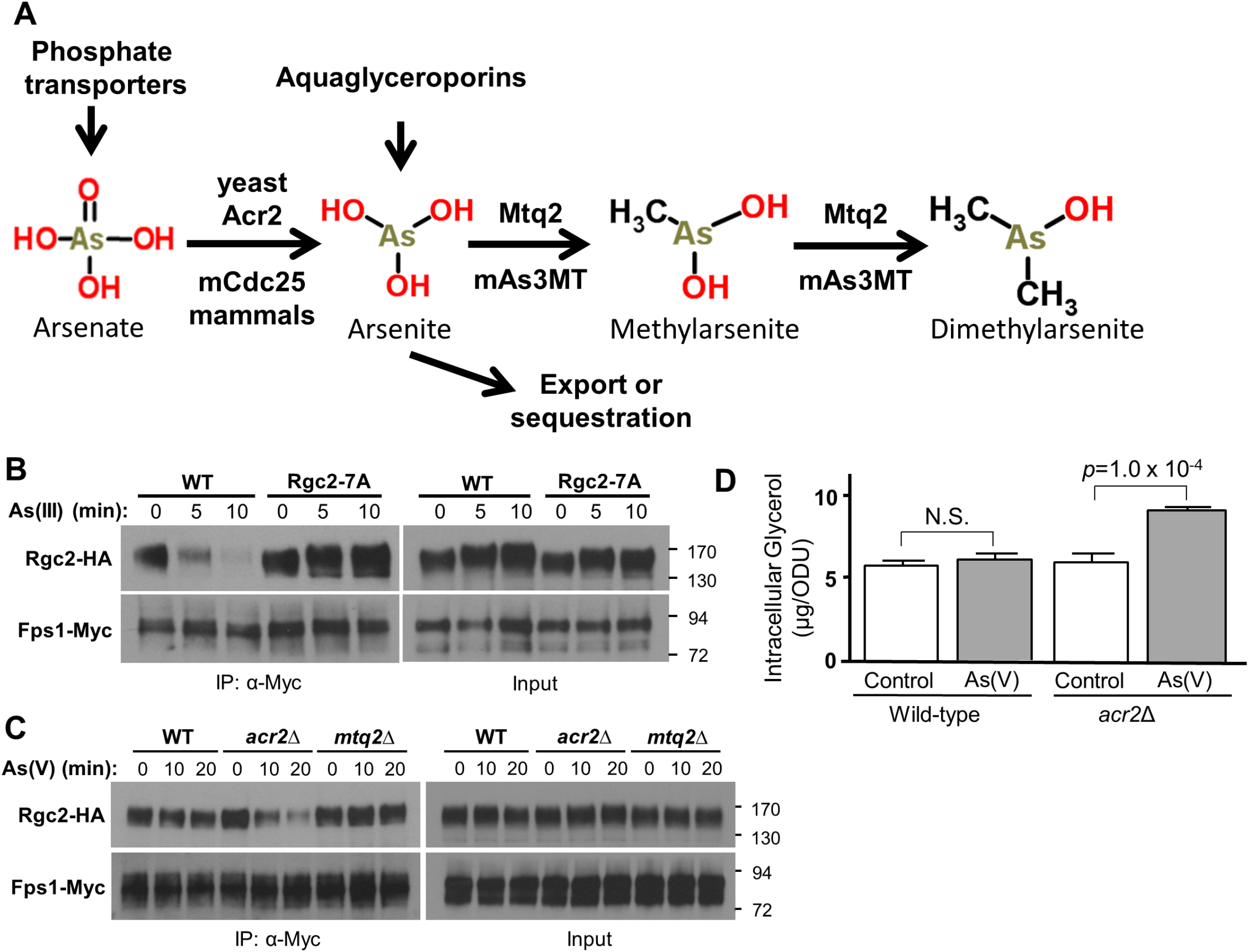
Regulation of glycerol channel Fps1 in response to arsenate and arsenite. **A** Metabolism of arsenicals in yeast and mammals. Pentavalent arsenate [As(V)] enters the cell through phosphate transporters and is reduced to trivalent arsenite [As(III)], a required step for export from the cell or sequestration in the vacuole. As(III) enters the cell mainly through aquaglyceroporins and is methylated to methylarsenite [MAs(III)], which can be further methylated to dimethylarsenite [DMA(III)]. The responsible enzymes are noted for yeast and mammals. **B** Closure of Fps1 in response to As(III) exposure requires Hog1 phosphorylation sites on its regulator, Rgc2. Wild-type Rgc2-HA (p3151), or a mutant form lacking seven Hog1 phosphorylation sites (Rgc2-7A-HA; p3155), was co-expressed with Fps1-Myc (p3121) in an *rgc1*Δ *rgc2*Δ strain (DL3207). Fps1 closure was detected by dissociation of Rgc2 in co-IP after treatment with 1mM As(III) for the indicated times. Anti-Myc IPs were separated by SDS-PAGE and subjected to immunoblot analysis. Molecular mass markers (in kDa) are shown on the right. **C** As(V) exposure induces closure of Fps1 in an *acr2*Δ mutant. Rgc2-HA (p3151) and Fps1-Myc (p3121) were co-expressed in a wild-type strain (DL3187), an *acr2*Δ mutant (DL4341), or an *mtq2*Δ mutant (DL4313). The strains were treated with 3 mM As(V) for the indicated times and processed for co-IP, as above. **D** An *acr2*Δ mutant accumulates glycerol in response to As(V) treatment. A wild-type strain (DL3187) and an *acr2*Δ mutant (DL4341) were grown in logarithmic phase with, or without, 3 mM As(V) for two hours prior to measurement of intracellular glycerol levels. Each value is the mean and standard deviation from three independent cultures. Pair-wise *p*-values for As(V)-treated and untreated samples were calculated using student t-test. N.S.; Not significant.

To address the question of how the cell differentially regulates Fps1 in response to arsenicals, we used co-immuneprecipitation (co-IP) to assess Hog1-driven dissociation of Rgc2 from Fps1 as an assay for Fps1 closure (Lee et al., 2013; Lee and Levin, 2018). In response to hyper-osmotic stress, Hog1 phosphorylates Rgc2 on at least seven Ser/Thr residues, which triggers its dissociation from Fps1 and consequent channel closure (Lee et al., 2013). Mutation of these residues to Ala (Rgc2-7A) blocks Rgc2 dissociation in response to hyper-osmotic stress. We found that Rgc2-7A was similarly stabilized on Fps1 in response to As(III) activation of Hog1 (Figure 1B), suggesting that Hog1 phosphorylates the same constellation of residues on Rgc2 in response to both As(III) treatment and hyper-osmotic stress. We next asked if blocking metabolism of As(V) to As(III), or As(III) to MAs(III), might influence Hog1-driven closure of Fps1. We found that blocking reduction of As(V) to As(III) with an *acr2*Δ mutation allowed Hog1 to close Fps1 in response to As(V) treatment (Figure 1C) and to accumulate glycerol to a modest degree (Figure 1D). This result revealed that the production of As(III) is important for blocking Fps1 closure in response to As(V) treatment and raised the possibility that modification of at least one arsenylation target prevents Hog1 from closing Fps1. In contrast to this, blocking conversion of As(III) to MAs(III) with an *mtq2*Δ mutation did not allow Fps1 closure in response to As(V) treatment (Figure 1C) despite activation of Hog1 in this mutant (Lee and Levin, 2018), suggesting that MAs(III) production does not play a role in blocking As(V)-activated Hog1 from closing this channel.

### The one-step metabolism model

The results above reveal that As(III) produced from exposure to As(V) prevents active Hog1 from closing Fps1. However, this finding is somewhat counterintuitive, because Hog1 activated in response to As(III) treatment drives Fps1 closure. Because As(V) is metabolized to As(III), which is metabolized to MAs(III) (Figure 1A), the cell is faced with the problem of identifying the primary threat and responding appropriately. We propose that the cell solves this problem by allowing arsenicals to be metabolized through only a single step. That is, As(V) is metabolized to As(III), but not further; whereas when cells are exposed to As(III), it is metabolized to MAs(III). This would result in the modification of a set of cysteine thiols by As(III) in response to either As(V) or As(III) challenge and an additional set of cysteine thiols modified specifically by MAs(III) only in response to As(III) challenge (Figure 2A). This arrangement is possible because of the atypical nature of the arsenic metabolism pathway. In contrast to most metabolic pathways in which each enzyme has a continuous supply of substrate to drive subsequent steps, the cell minimizes the intracellular concentration of As(III) produced from As(V) through its rapid extrusion from the cell (through Acr3 and Fps1; Wysocki et al., 2001; Maciaszczyk-Dziubinska et al., 2012), and sequestration in the vacuole after its conjugation to glutathione (Maciaszczyk-Dziubinska et al., 2011, 2012). Cysteine residues that are targets of As(III) modification may also be modified by MAs(III) (Figure 2A, dotted arrow).

**Figure 2.**
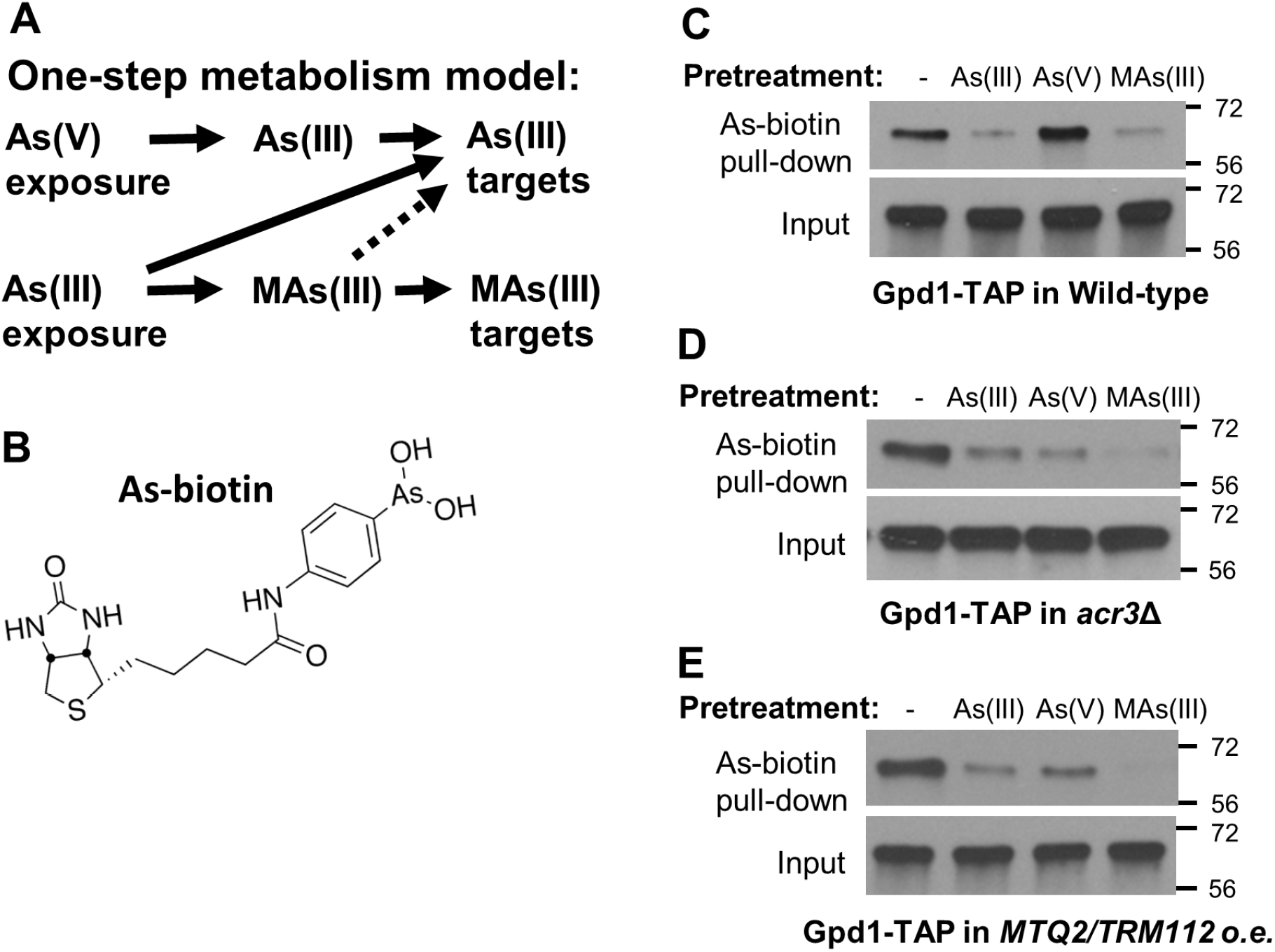
The one step metabolism model. **A** We propose that the two major forms of arsenic in the environment are each metabolized by only one step. As(V) exposure results in the production of As(III) and consequent arsenylation of a set of cysteine thiols in target proteins. As(III) exposure results in the production of MAs(III) and methylarsenylation of a set of cysteine thiols in addition to those modified by As(III). MAs(III) produced from As(III) exposure may also modify cysteine residues targeted by As(III) (dotted arrow). **B** The structure of the As-biotin conjugate used. **C** In vivo binding of As-biotin to Gpd1-TAP is diminished by pretreatment with As(III) or MAs(III), but not As(V). A wild-type strain (DL3187) transformed with a plasmid expressing Gpd1-TAP under the inducible control of the *GAL1* promoter (p3467) was pre-treated for 10 min with 1 mM As(III), 3 mM As(V), or 0.5 mM MAs(III) prior to a 10- min treatment with 10 μM As-biotin. Extracts were subjected to affinity pull-down with streptavidin agarose (SA) beads prior to SDS-PAGE and immunoblot analysis for Gpd1- TAP. Molecular mass markers (in kDa) are on the right. **D** In vivo binding of As-biotin to Gpd1-TAP is diminished by forced production of MAs(III) from As(V) pretreatment in an *acr3*Δ mutant. An *acr3*Δ mutant (DL4287) transformed with a plasmid expressing Gpd1-TAP (p3467) was treated as above. E. In vivo binding of As-biotin to Gpd1-TAP is diminished by forced production of MAs(III) from As(V) pretreatment by overexpression of the As(III) methyltransferase. A wild-type strain (DL3187) was co-transformed with a plasmid expressing Gpd1-TAP (p3467) and plasmids over-expressing (o.e.) both subunits of the As(III) methyltransferase (Mtq2 [p3629] and Trm112 [p3460]) under the repressible control of the *MET25* promoter. The strain was treated as above.

However, the key requirements of this model are: 1) the existence of Cys residues that are modified exclusively by MAs(III), and 2) that MAs(III) is not produced by As(V) treatment. Modification of these Cys residues would, therefore, signal an As(III) challenge.

We have developed evidence that supports our contention that As(V) challenge does not result in the production of physiologically significant levels of MAs(III). First, using an arsenic-biotin probe (As-biotin; Figure 2B) in vivo, coupled with pull-down of covalently bound proteins from lysates with streptavidin beads, we have identified specific cysteine residues in target proteins that are modified by MAs(III), but not by As(III). For example, Cys306 in Gpd1 is a MAs(III)-specific target, modification of which results in the inhibition of Gpd1 catalytic activity (Lee and Levin, 2019). Pre-treatment of an *mtq2*Δ mutant with MAs(III), but not with As(III), diminishes As-biotin binding to Gpd1, establishing MAs(III) as the natural arsenical that modifies Gpd1. On the other hand, As-biotin binding to Gpd1 was diminished in wild-type cells pre-treated with either As(III) or MAs(III) (Figure 2C; and Lee and Levin, 2019), because As(III) is metabolized to MAs(III) in this setting. In contrast to this, pre-treatment of wild-type cells with As(V) did not diminish As-biotin binding to Gpd1 (Figure 2C), suggesting that As(V) is not metabolized to physiologically significant levels of MAs(III).

We extended this analysis with two approaches to force MAs(III) production from As(V). First, loss of *ACR3*, the gene that encodes the As(III) efflux pump, results in the accumulation of intracellular As(III) (Maciaszczyk-Dziubinska et al., 2011). We found that As(V) pre-treatment of an *acr3*Δ mutant diminished As-biotin binding to Gpd1 (Figure 2D), supporting the conclusion that As(V) is not normally metabolized to MAs(III), but it can be forced by mass action. The second approach to drive MAs(III) production was by co-overexpression of *MTQ2* and *TRM112*, which encode the dimeric As(III) methyltransferase (Lee and Levin, 2018). As(V) pre-treatment diminished As-biotin binding to Gpd1 in this setting as well (Figure 2E), suggesting that this enzyme may be rate-limiting in the metabolism of As(V). Taken together, these results support the conclusion that As(V) is normally metabolized to As(III), but not to MAs(III). We further conclude that under conditions of As(V) exposure, both free intracellular As(III) and As(III) methyltransferase may be limited.

### Hog1 is modified by As(III)

The results presented above suggest that arsenylation of a cellular target prevents As(V)-activated Hog1 from closing Fps1. Therefore, we asked if Hog1 itself might be arsenylated. We found that As-biotin binds to Hog1 in vivo (Figure 3A). Therefore, we explored the possibility that Hog1 modification by As(III) is responsible for its failure to close Fps1 in response to As(V) challenge. There are three cysteine residues within the Hog1 catalytic domain that are conserved with mammalian p38 SAPK (Cys38, Cys156, and Cys205). We mutated each of these, individually and in combination, to serine residues to prevent their modification by arsenicals. The only single mutant to display hyper-sensitivity to growth inhibition by As(III) was *hog1-C156S* (Figure 3B). Both double mutants with *C156S* displayed further sensitivity, as did the triple mutant (*hog1-3C/S*). In contrast to this, none of these mutant forms of Hog1 resulted in growth sensitivity under hyper-osmotic stress conditions (i.e. 1M sorbitol).

**Figure 3.**
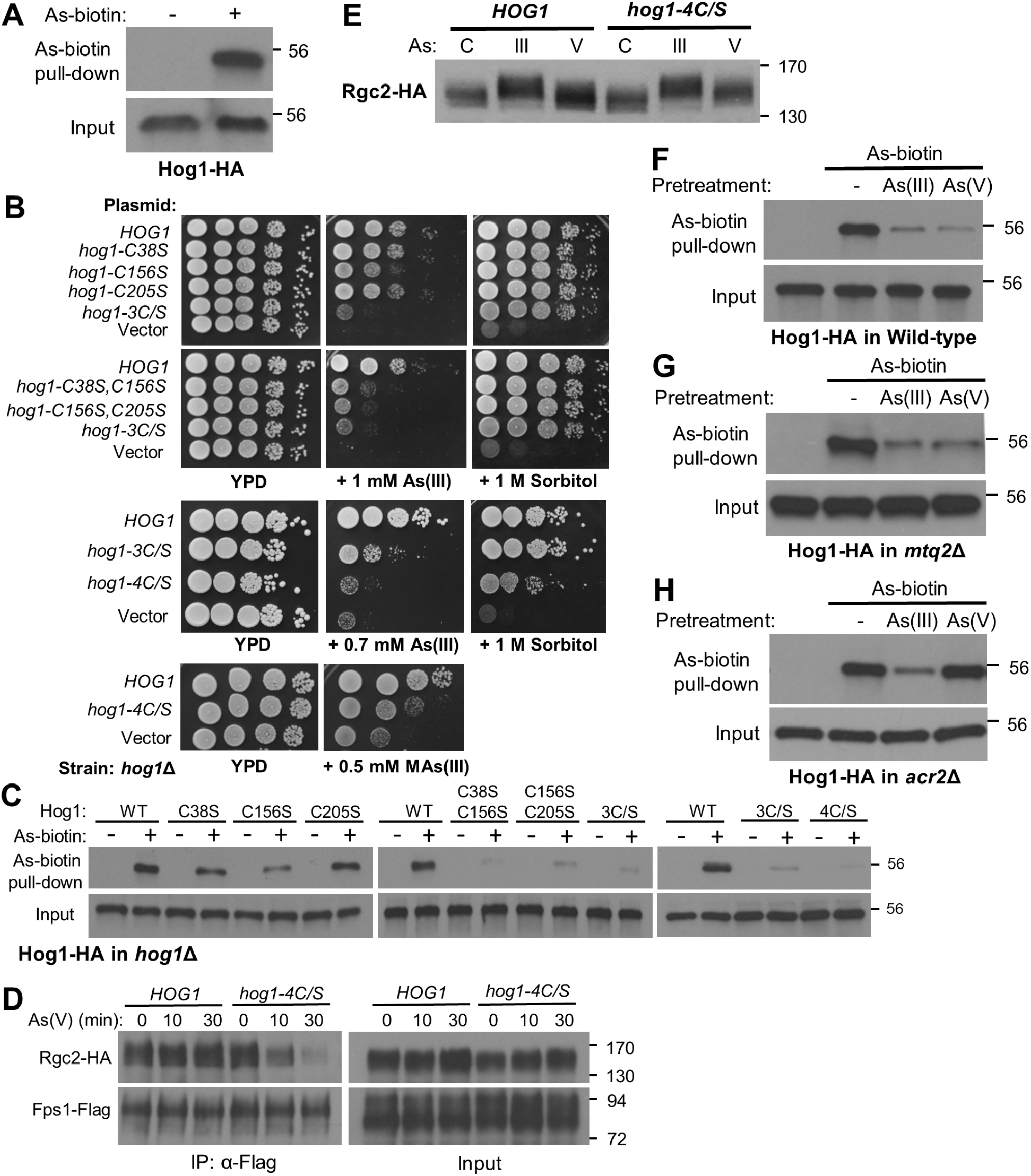
Hog1 is arsenylated. **A** As-biotin binds to Hog1-HA in vivo. A wild-type strain (DL3187) transformed with a multi-copy plasmid expressing Hog1-HA (p3225) was treated (+) or not (-) with 10 μM As-biotin for 10 min. Extracts were subjected to affinity pull-down with SA beads prior to SDS-PAGE and immunoblot analysis for Hog1-HA. Molecular mass marker (in kDa) is on the right. **B** Mutations in *HOG1* at cysteine residues cause As(III) hyper-sensitivity. Cultures of a *hog1*Δ strain (DL3158) transformed with centromeric plasmids expressing the indicated *HOG1* allele were spotted onto YPD, YPD plus the indicated concentration of As(III) or MAs(III), or YPD plus 1M sorbitol, at serial 10-fold dilutions (from left to right) and incubated at 30°C for 3 days. Plasmids carried *HOG1* (p3090), *hog1-C38S* (p3577), *hog1-C156S* (p3578), *hog1-C205S* (p3579), *hog1-C38S, C156S* (p3580), *hog1-C156S, C205*S (p3581), *hog1-C38S, C156S, C205S* (*hog1-3C/S*; p3582), *hog1-C38S, C156S, C161S, C205S* (*hog1-4C/S*; p3583), or vector (p3090). **C** Four cysteine residues in Hog1 are bound by As-biotin. Mutant forms of Hog1-HA were tested for As-biotin binding using the same strains as in “C” after treatment with 10 μM As-biotin for 10 min. **D** The *hog1-4C/S* mutant drives Fps1 closure in response to As(V) exposure. A *hog1*Δ strain (DL3158) co-transformed with plasmids expressing Rgc2-HA (p3471), Fps1-Flag (p2492), and either wild-type Hog1 (p3090) or Hog1-4C/S (p3583) was treated with 3 mM As(V) for the indicated times prior to processing for co-IP with anti-Flag antibodies. **E** As(V) treatment induces Rgc2 phosphorylation in the *hog1-4C/S* mutant. A *hog1*Δ strain (DL3158) co-transformed with plasmids expressing Rgc2-HA (p3182) and either wild-type Hog1 (p3090) or Hog1-4C/S (p3583) was exposed to 1 mM As(III) for 10 min, 3 mM As(V) for 20 min, or untreated (C) and extracts were subjected to SDS-PAGE on a 7.5% gel to resolve phosphorylated forms of Rgc2-HA. **F-H** As-biotin pull-down of Hog1 from wild-type cells, an *mtq2*Δ mutant, or an *acr2*Δ mutant with pre-treated with either As(III) or As(V). Wild-type cells (F; DL3187), an *mtq2*Δ mutant (G; DL4313), or an *acr3*Δ mutant (H; DL4341) expressing Hog1-HA (p3225) were pre-treated with 1 mM As(III) or 3 mM As(V) for 20 min. prior to treatment with 10 μM As-biotin for 10 min. Extracts were subjected to affinity pull-down with SA beads prior to SDS-PAGE and immunoblot analysis for Hog1-HA.

These results mirrored our findings of As-biotin binding to Hog1, which was somewhat diminished in the Hog1-C156S form and further diminished by the double and triple mutations (Figure 3C). However, some As-biotin binding to Hog1 remained even in the Hog1-3C/S mutant form. Guerra-Moreno et al. (2019) reported that Hog1 becomes modified on three cysteine residues -- Cys156, Cys161, and Cys205 in response to As(III) treatment. Because we had not included Cys161 in our initial analysis, we added the Cys161Ser mutation to the *hog1-3C/S* allele, yielding *hog1-4C/S*. The quadruple mutant was even more sensitive to growth inhibition by As(III) than was the *hog1-3C/S* mutant (roughly equivalent to the vector control), but it also displayed slight growth sensitivity under hyper-osmotic stress conditions (Figure 3B). Significantly, As-biotin binding to Hog1-4C/S was further diminished as compared with the triple mutant form (Figure 3C). We conclude that these four cysteine residues are important for proper Hog1 response to As(III) challenge and are modified by an arsenical.

We next tested the *hog1-4C/S* mutant for its ability to close Fps1 in response to As(V) challenge. Intriguingly, this mutant responded to As(V) treatment by inducing eviction of Rgc2 from Fps1 (Figure 3D), suggesting that arsenic modification of cysteine thiols in Hog1 is responsible for blocking its ability to close Fps1. These findings were further supported by the detection of an Rgc2 hyper-phosphorylation band-shift in response to As(V) treatment only in the *hog1-4C/S* mutant (Figure 3E). Although the band-shift was not as great as that observed in response to As(III) treatment, this was as expected because only a fraction of the Rgc2 phosphorylation induced in response to As(III) exposure is catalyzed by Hog1 (Beese et al., 2009; Lee et al., 2013). The *hog1-4C/S* mutant closed Fps1 normally in response to As(III) treatment (Supplemental Figure S1), indicating that the hyper-sensitivity of this mutant to As(III) is caused by some other deficiency in its response.

Because we have found that some cysteine residues in proteins are specifically targeted by MAs(III) (Lee and Levin, 2018; 2019), we examined whether Hog1 is modified by As(III), or by MAs(III). Pre-treatment of wild-type cells with either As(III) or As(V) diminished As-biotin binding to Hog1 (Figure 3F). Similar results were obtained by pre-treatment of an *mtq2*Δ mutant with either arsenical, revealing that Hog1 is modified by As(III) in response to either challenge, without a requirement for the production of MAs(III) (Figure 3G). In contrast to this, pre-treatment of an *acr2*Δ mutant with As(V) failed to diminish As-biotin binding to Hog1 (Figure 3H), supporting the conclusions that As(V) must be reduced to As(III) to modify Hog1 and that Acr2 is the major (or only) arsenate reductase in yeast. We also found that pre-treatment with MAs(III) blocked As-biotin binding to Hog1 (Supplemental Figure S2), but it is not clear if this finding has biological significance in the context of an As(III) challenge.

In summary, we conclude that Hog1 is modified by As(III) on four Cys residues in response to either As(III) or As(V) treatment and that these modifications prevent As(V)- activated Hog1 from phosphorylating Rgc2 and closing Fps1. To assess the biological impact of preventing Fps1 closure in response to As(V) exposure, we compared the growth of a *hog1*Δ strain expressing *HOG1*, *hog1-4C/S*, or empty vector in the presence of As(V). Although a spot growth plate did not show a growth deficiency of the *hog1- 4C/S* mutant in the presence of 5 mM As(V) (Supplemental Figure S3), we detected a reduced growth rate for this mutant in liquid culture. In the absence of treatment, all three strains grew comparably on selective medium (with a doubling time of 105 min.).

Treatment with 1 mM As(V) impaired the growth of a *hog1*Δ mutant to approximately the same degree as wild-type (doubling times of 146 min. vs. 143 min. respectively), but growth of the *hog1-4C/S* mutant was impaired reproducibly to a greater degree (doubling time of 157 min.), revealing that some function of the Hog1-4C/S form is responsible for the reduced growth rate. We suggest that this function is the closure of Fps1 in an inappropriate setting, which results in both increased glycerol and As(III) accumulation. Finally, both the *hog1-4C/S* mutant and the *hog1*Δ mutant displayed increased sensitivity to MAs(III) treatment relative to wild-type (Figure 3B), consistent with our previous finding that MAs(III) treatment activates Hog1 (Lee and Levin, 2018).

### Acr3 is a critical regulator of Fps1

We have shown above that modification of Hog1 by As(III) in response to As(V) treatment blocks Fps1 closure. However, Hog1 activated in response to As(III) treatment is similarly modified, yet it induces Fps1 closure. The one-step metabolism model predicts that, in response to As(III) challenge, a target of MAs(III) modification would allow arsenylated Hog1 to overcome this block. In this section, we demonstrate that the plasma membrane As(III) efflux pump, Acr3, is a key regulator of Fps1 closure whose modification by MAs(III) relieves the block imposed by arsenylation of Hog1.

We found in a screen of null mutants in genes that are important for the arsenic response that As(V) treatment drives Fps1 closure in an *acr3*Δ mutant (Figure 4A), suggesting that Acr3 may be a regulator of Fps1 activity. Although we showed above that loss of *ACR3* results in the production of MAs(III) from As(V) (Figure 2D), thereby mimicking an As(III) challenge, we demonstrate below that Acr3 also regulates Fps1 directly. Because Acr3 is a plasma membrane protein, we first tested by co-IP whether Acr3 is a component of the Fps1 complex. We immuneprecipitated epitope-tagged Acr3 (Acr3-HA) from extracts with differentially tagged Fps1 (Fps1-Myc) and found that Fps1- Myc co-precipitated with Acr3-HA (Figure 4B). This association did not appear to be altered in response to either As(V) or As(III) treatment (Supplemental Figure S4). We also explored the interaction between these two plasma membrane proteins using Bimolecular Fluorescence Complementation (BiFC). This method is used to visualize in vivo interactions of proteins tagged with two halves of a cyan fluorescent protein (CFPN or CFPC), which are united to restore fluorescence signal (Lipatova et al., 2012). We found that a CFPC-Fps1 chimera associates with an Acr3-CFPN chimera at the plasma membrane in wild-type cells (Supplemental Figure S5). Importantly, this interaction was retained in a strain in which genes that encode other known components of the Fps1 complex (i.e. *HOG1*, *RGC1* and *RGC2*) were deleted (Figure 4C), suggesting that the interaction between Acr3 and Fps1 may be direct. We also asked if Acr3 could associate with other components of the Fps1 complex. We detected an Acr3-CFPN interaction with both CFPC-Rgc2 and CFPC-Hog1 in wild-type cells, but the Acr3- CFPN/CFPC-Hog1 interaction was not detected in the absence of Fps1 (Supplemental Figure S5), suggesting that this association is indirect. Finally, we found that the Acr3- CFPN/CFPC-Rgc2 interaction was retained in the absence of Fps1 (Figure 4D), suggesting that this interaction may, like the Acr3/Fps1 interaction, be direct.

**Figure 4.**
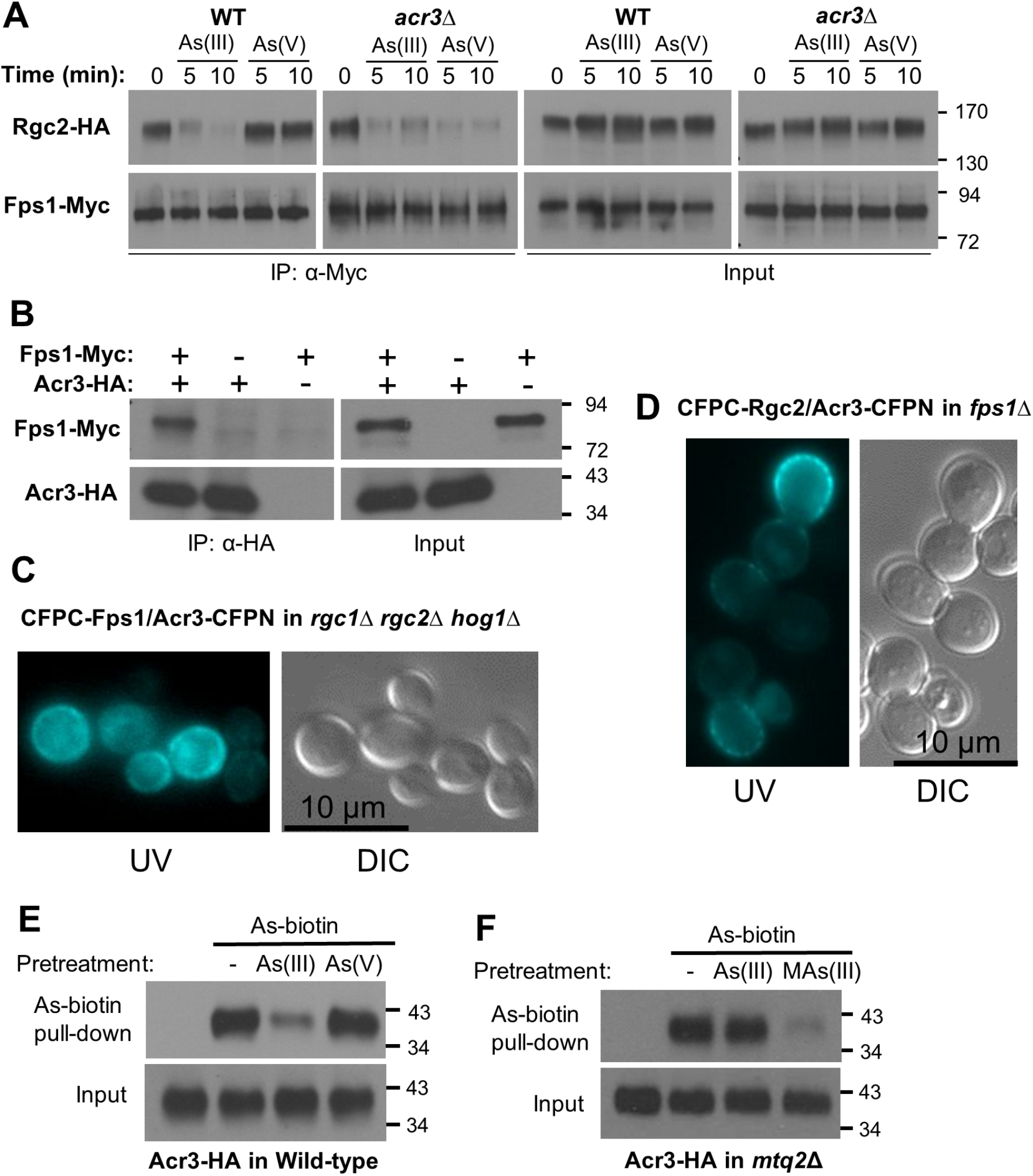
Acr3 is a component of the Fps1 complex that regulates Fps1 in response to arsenicals. **A** As(V) treatment induces Fps1 closure in an *acr3*Δ mutant. Rgc2-HA and Fps1-Myc were co-expressed (from p3151 and p3121, respectively) in a wild-type strain (DL3187), or an *acr3*Δ mutant (DL4287). The strains were treated with 1 mM As(III) or 3 mM As(V) for the indicated times and processed for co-IP of Rgc2 with Fps1 using anti-Myc antibodies, as in Figure 1. **B** Acr3 associates with Fps1. A wild-type strain (DL3187) co-expressing Fps1-Myc (p3121) and Acr3-HA (p3470), or expressing only one protein was subjected to co-IP analysis using anti-HA antibodies. **C and D** Bimolecular Fluorescence Complementation (BiFC) analysis of Acr3 with known components of the Fps1 complex. An *rgc1*Δ *rgc2*Δ *hog1*Δ strain (C; DL3219) co- expressing CFPC-Fps1 (p3216) and Acr3-CFPN (p3585) was visualized under a UV light source to reveal fluorescence complementation, or under visible light (DIC). An *fps1*Δ strain (D; DL3226) co-expressing CFPC-Rgc2 (p3584) and Acr3-CFPN (p3585) was visualized as above. Representative micrographs are shown. **E** Pre-treatment of wild-type cells with As(III), but not As(V), blocked As-biotin binding to Acr3-HA. Wild-type strain (DL3187), expressing Acr3-HA (p3470) was pre-treated with 1 mM As(III) or 3 mM As(V) for 20 min. prior to treatment with 10 μM As-biotin for 10 min. Extracts were subjected to affinity pull-down with SA beads prior to SDS-PAGE and immunoblot analysis for Acr3-HA. **F** Acr3 is a MAs(III) target. An *mtq2*Δ mutant (DL4313) expressing Acr3-HA (p3470) was pre-treated with 1 mM As(III) or 0.5 mM MAs(III) for 20 min. prior to treatment with 10 μM As-biotin for 10 min. and subjected to affinity pull-down, as above. Molecular mass markers (in kDa) are on the right.

We next asked if Acr3 is a target of arsenic binding using the As-biotin probe. We detected Acr3-HA in streptavidin bead pull-downs of extracts from cells treated with As-biotin and this binding was diminished in wild-type cells by pre-treatment with As(III), but not with As(V) (Figure 4E), suggesting that Acr3 is modified by MAs(III). We tested this conclusion directly by repeating this experiment in an *mtq2*Δ mutant to eliminate the complication of As(III) metabolism to MAs(III). Pre-treatment of this mutant with As(III) failed to diminish As-biotin binding to Acr3-HA, indicating that As-biotin does not detect any As(III)-binding sites in Acr3 (Figure 4F). However, pre-treatment with MAs(III) blocked As-biotin binding, revealing the presence of at least one MAs(III)-binding site in Acr3. Moreover, the failure of As(V) pre-treatment of wild-type cells to block As-biotin binding to Acr3 provides further support for the one-step metabolism model.

Members of the Acr3 family are multi-pass transmembrane proteins and fungal versions of this transporter differ most notably from bacterial and plant forms by an extended cytoplasmic loop #4 (Maciaszczyk-Dziubinska et al. 2014), which possesses three cysteine residues (Cys316, Cys318, and Cys333; Figure 5A). A form of Acr3 that bears a 28-amino acid deletion within this loop was shown to be fully functional with regard to As(III) transport (Wawrzycka et al., 2017). Therefore, we asked if this loop is important for the interactions of Acr3 with Fps1 and Rgc2. We re-created this deletion form of Acr3 (Acr3-Δ307-334), as well as an Acr3-Δ307-334-CFPN fusion for use in BiFC. The *ACR3-Δ307-334* allele was able to complement fully the extreme As(III) sensitivity of an *acr3*Δ mutant (Figure 5B), confirming that it is functional. In fact, this mutant provided slightly greater As(III) tolerance than the wild-type allele. The Acr3- Δ307-334-CFPN protein was able to interact at the plasma membrane with CFPC-Fps1 in an *rgc1*Δ *rgc2*Δ *acr3*Δ strain (Figure 6A), indicating that cytoplasmic loop #4 is not required for this interaction. Intriguingly, however, this mutant protein was not able to interact with CPFC-Rgc2 in an *fps1*Δ *acr3*Δ strain (Figure 6B), revealing that cytoplasmic loop #4 is required for the Acr3 interaction with Rgc2 and suggesting that this loop may interact directly with Rgc2. An updated model of the Fps1 complex that includes Acr3 is shown in Figure 6C. This model takes into account our previous demonstrations that Hog1 binds to a site on the N-terminal extension of Fps1 and that Rgc2 (and Rgc1) binds to a site on the C-terminal Fps1 extension (Lee et al., 2013).

**Figure 5.**
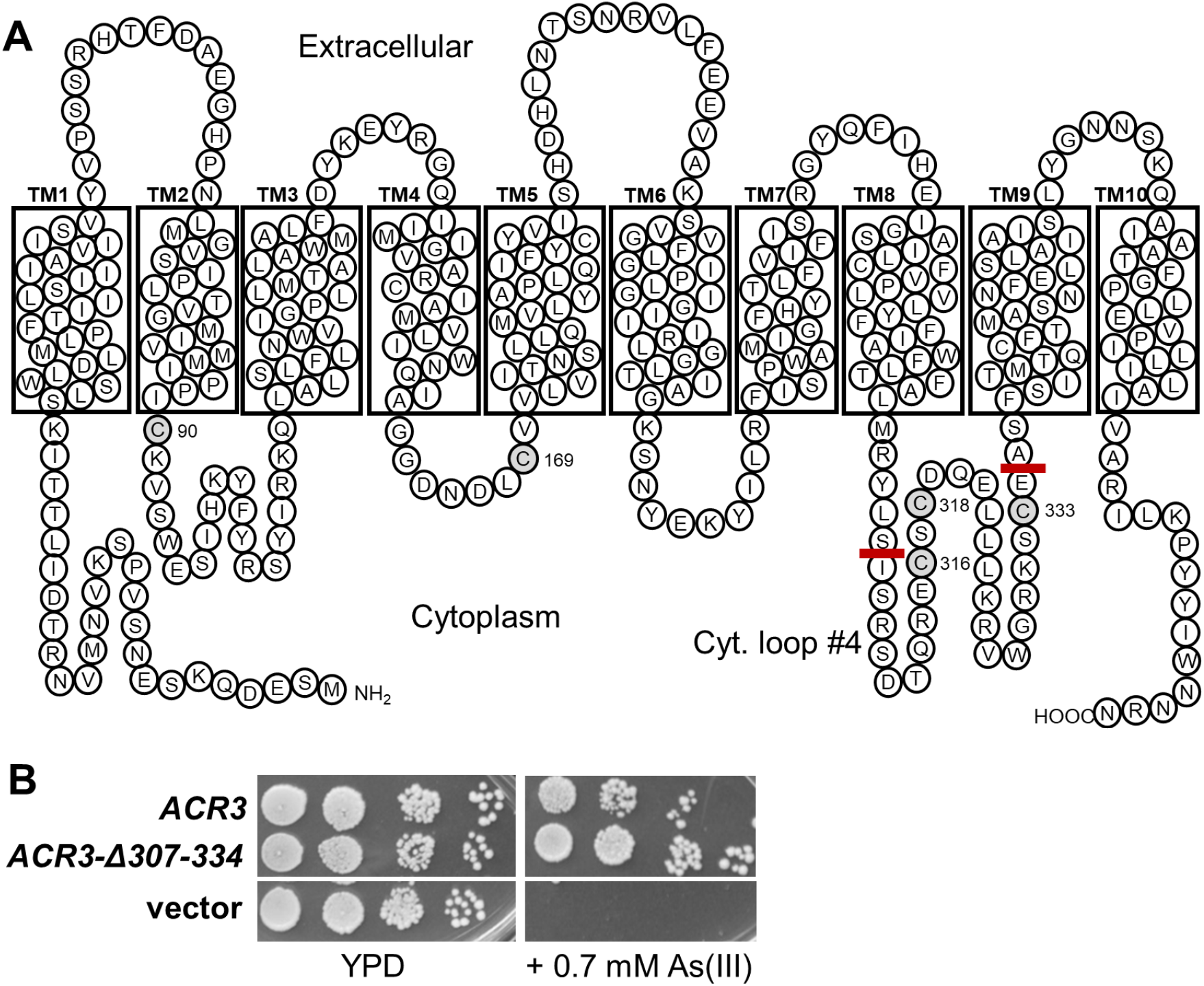
Arsenite efflux pump Acr3. **A** Acr3 membrane topology (after Maciaszczyk-Dziubinska et al., 2014), showing cytoplasmic cysteine residues (marked in gray) and the cytoplasmic loop #4 (marked with red bars) that is unique to fungal forms of Acr3 (Wawrzycka et al., 2017). **B** A deletion mutant that removes 28 residues within cytoplasmic loop #4 of Acr3 complements the As(III) sensitivity of an *acr3*Δ mutant better than does wild-type *ACR3*. Cultures of an *acr3*Δ mutant (DL4287) transformed with centromeric plasmids expressing *ACR3* (p3587) or *ACR3-Δ307-334* (p3588) were spotted onto YPD, or YPD plus 0.7 mM As(III), at serial 10-fold dilutions (from left to right) and incubated at 30°C for 3 days.

**Figure 6.**
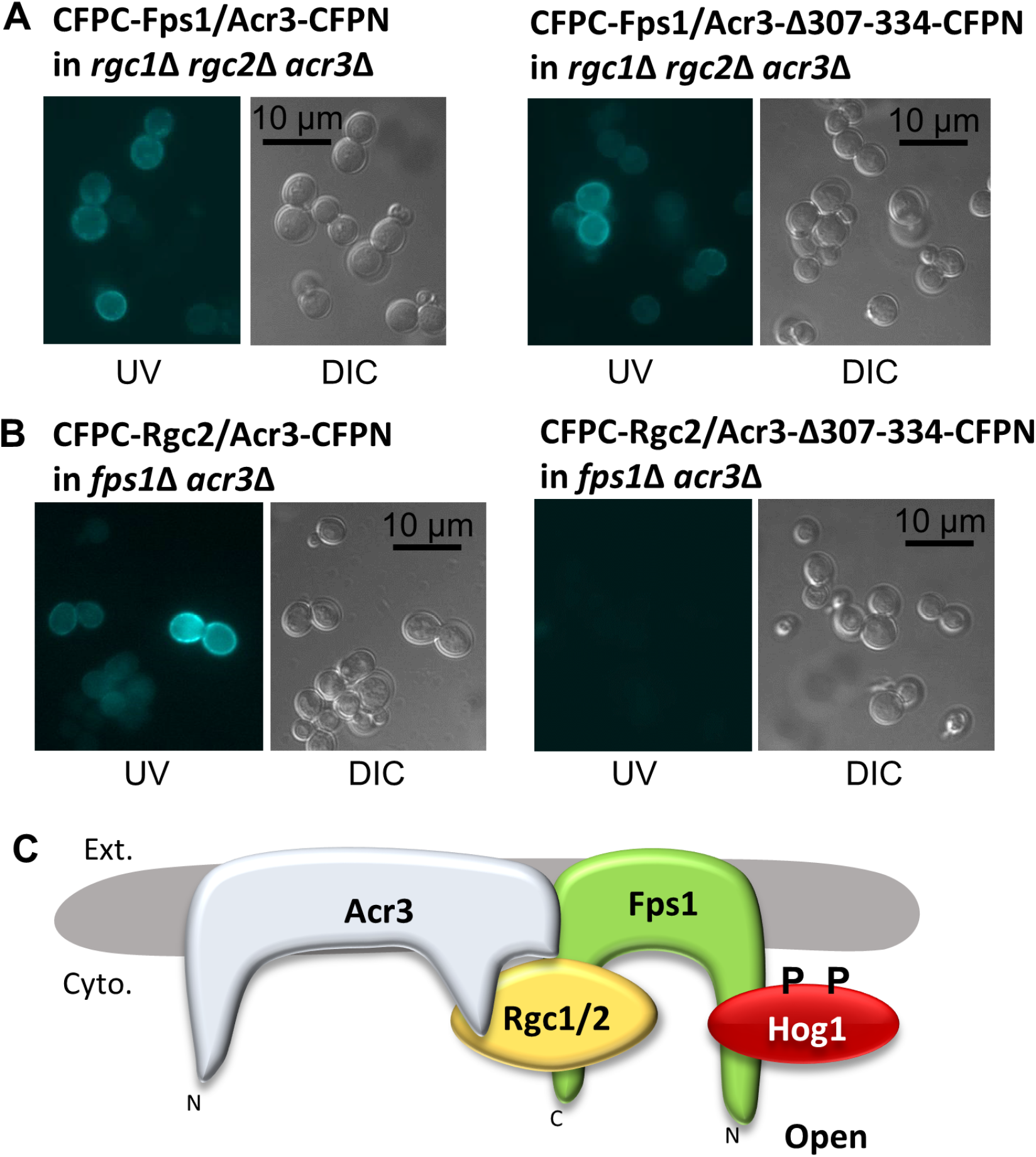
Acr3 cytoplasmic loop #4 is required for its interaction with Rgc2. **A and B** BiFC showing interactions between Acr3 or Acr3-Δ307-334, and Fps1 or Rgc2. A. An *rgc1*Δ *rgc2*Δ *acr3*Δ strain (DL4273) expressing CPFC-Fps1 (p3216) and either Acr3-CFPN (p3585) or Acr3-Δ307-334-CFPN (p3586) was visualized under a UV light source to reveal fluorescence complementation, or under visible light (DIC). B. An *fps1*Δ *acr3*Δ strain (DL4271) expressing CFPC-Rgc2 (p3584) and either Acr3-CFPN or Acr3- Δ307-334-CFPN was visualized as above. Representative micrographs are shown. **C** Model of the Fps1 complex showing association of phospho-Hog1 and Rgc2 (and Rgc1) with the N-terminal and C-terminal extensions of Fps1, respectively. The model also shows association of Acr3 with Fps1 and the Acr3 cytoplasmic loop #4 with Rgc2.

We next asked how the *ACR3-Δ307-334* allele behaves with regard to the regulation of Fps1. Like the *acr3*Δ mutant, this mutant allowed Fps1 closure in response to As(V) treatment (Figure 7A), indicating that cytoplasmic loop #4 normally prevents Hog1 activated in response to As(V) from closing Fps1, and suggesting that it may protect Rgc2 from phosphorylation by arsenylated Hog1. Therefore, to extend our finding that Acr3 is modified by MAs(III), we explored the possibility that the three cysteine residues within the Acr3 cytoplasmic loop #4 might be the targets of MAs(III) in response to As(III) treatment, which would disrupt the interaction of loop #4 with Rgc2. We mutated these cysteine residues to serine, (Cys333 alone and together with a Cys316 Cys318 double mutation) and tested these mutant forms of Acr3-HA for their ability to bind As-biotin. Figure 7B shows that As-biotin bound less well to either the Acr3-C333S or Acr3-C316/318/S (Acr3-2C/S) forms than to wild-type Acr3 and that As-biotin did not bind detectably to the Acr3-C316/318/333S(Acr3-3C/S) form, revealing that all three of these cysteine residues are methylarsenylated.

**Figure 7.**
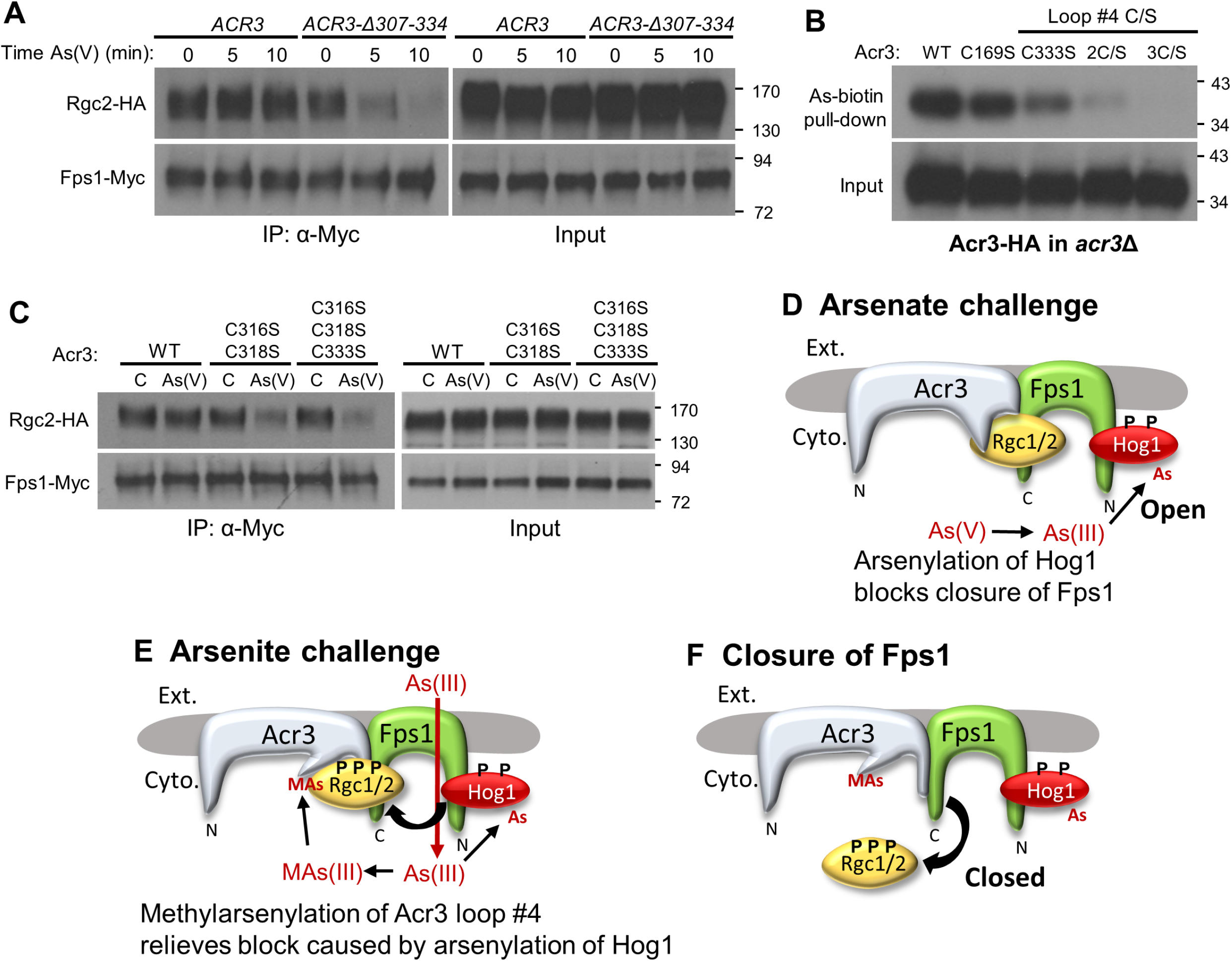
Cysteine residues in the Acr3 cytoplasmic loop #4 are modified by MAs(III) and are important for the regulation of Fps1. **A** Effect of Acr3 cytoplasmic loop #4 deletion on Fps1 closure in response to As(V) treatment. An *acr3*Δ strain (DL4287) co-transformed with plasmids expressing Fps1- Myc (p3121), Rgc2-HA (p3151) and either *ACR3* (p3587) or *ACR3-Δ307-334* (p3588) were treated with 3 mM As(V) for the indicated times and processed for co-IP of Rgc2 with Fps1 using anti-Myc antibodies. **B** The three cysteine residues within Acr3 loop #4 account for its As-biotin binding. An *acr3*Δ strain (DL4287) expressing the indicated form of Acr3 was treated with 10 µM As- biotin for 10 min. Extracts were subjected to affinity pull-down with SA beads prior to SDS-PAGE and immunoblot analysis for Acr3-HA. Plasmids expressed one of the following *ACR3* alleles: *ACR3-HA* (p3470), *acr3-C169S-HA* (p3591), *acr3-C333S-HA* (p3592), *acr3-C316S, C318S-HA* (2C/S; p3593), or *acr3-C316S, C318S, C333S-HA* (3C/S; p3594). **C** Cys-to-Ser mutations within Acr3 loop #4 allow closure of Fps1 in response to As(V) treatment. An *acr3*Δ strain (DL4287) was co-transformed with plasmids expressing Fps1-Myc (p3121), Rgc2-HA (p3151) and one of the following: *ACR3* (p3587), *acr3- C316S, C318S* (p3589), or *acr3-C316S, C318S, C333S* (p3590). Strains were either treated with 3 mM As(V) for 10 min, or untreated (C), and processed for co-IP of Rgc2 with Fps1. **D** As(V) challenge. As(V) is reduced to As(III), which modifies Hog1. Arsenylated Hog1 is prevented from phosphorylating Rgc2 (and presumably Rgc1), leaving Fps1 in the open state. **E** As(III) challenge. As(III) enters the cell through Fps1 and similarly modifies Hog1, but is also metabolized to MAs(III), which modifies Acr3 on cytoplasmic loop #4, which relieves the block of arsenylated Hog1 to phosphorylate Rgc1/2, **F** thereby inducing release of these regulators and consequent Fps1 closure.

Finally, we tested the ability of these mutant forms to regulate Fps1 properly in response to As(V) treatment. Like the *acr3*Δ mutant and the *ACR3-Δ307-334* mutant, the *ACR3-2C/S* and *ACR3-3C/S* alleles both allowed eviction of Rgc2 from Fps1 in response to As(V) treatment (Figure 7C), suggesting that these cysteine residues are important to protect Rgc2 from phosphorylation by arsenylated Hog1. However, the Acr3-3C/S form was still able to associate with Rgc2 in an *fps1*Δ *acr3*Δ strain as judged by BiFC (Supplemental Figure S5), suggesting that these residues are not the critical determinants for this association. We conclude from these data that arsenylation of Hog1 prevents it from closing Fps1 in response to As(V) challenge (Figure 7D), but this block is relieved in response to As(III) challenge by methylarsenylation of Acr3 on cytoplasmic loop #4 (Figure 7E), which allows Hog1 to phosphorylate Rgc2, causing its displacement from Fps1 and consequent closure of the channel (Figure 7F). This arrangement provides an elegant solution to the problem of mounting opposing responses to these related challenges.

## Discussion

The HOG pathway of yeast has been well characterized with regard to its regulation by hyper-osmotic stress (Saito and Posas, 2012). However, Hog1, the SAPK at the base of the HOG pathway, is also stimulated by a wide array of unrelated stress signals, including heat shock (Winkler et al., 2002), cold shock (Panadero et al., 2006), citric and acetic acid (Lawrence et al., 2004; Mollapour and Piper, 2006), oxidative stress (Bilsland et al., 2004), methylglyoxal (Aguilera et al., 2005), bacterial lipopolysaccharide (Marques et al., 2006), glucose starvation (Piao et al., 2012), curcumin (Azad et al., 2014), cadmium (Jiang et al., 2014), As(III) (Sotelo and Rodríguez-Gabriel, 2006; Thorsen et al., 2006) and As(V) (Lee and Levin, 2018).

Mammalian p38 SAPK is similarly activated by most of these stress-inducing agents (Ono and Han, 2000; de Nadel et al., 2002). This raises two important and related questions. First, do these various stresses activate the SAPK through a common pathway, or through alternative inputs? This is especially salient with regard to the HOG pathway because, until recently, the cell surface osmo-sensors at the head of this pathway were the only known inputs to Hog1. Second, how does an activated SAPK mount a specific response appropriate to the particular stress experienced? A problem of signal compression arises when a variety of stressors activate a common SAPK, but require distinct responses to be mobilized by the SAPK. We have begun to address these questions with an analysis of HOG pathway signaling in response to arsenic-induced stress.

### Single-step metabolism of arsenicals allows cells to distinguish between As(V) and As(III) challenges

In this study, we described the mechanism by which cells differentially regulate the glycerol channel Fps1 in response to Hog1 activation by either As(V) challenge or As(III) challenge. Hog1 closes Fps1 in response to As(III) treatment to restrict its entry through this adventitious port. On the other hand, Hog1 activated by As(V) exposure does not close Fps1, presumably to allow export of As(III) produced from As(V) metabolism (Lee and Levin, 2018). We presented evidence that cells distinguish between these stressors through single-step metabolism that results in the modification of cysteine thiols in target proteins by different trivalent arsenicals.

Specifically, when cells are exposed to As(V), it is reduced to As(III), which modifies a constellation of cysteine residues in various target proteins. On the other hand, when cells are exposed to As(III), it is metabolized to MAs(III), which modifies a different set of cysteine residues in target proteins in addition to those modified directly by As(III). Thus, the presence of intracellular MAs(III) signals an As(III) challenge, whereas the presence of intracellular As(III) without MAs(III) signals an As(V) challenge. We have shown previously that there exist specific targets of MAs(III) in response to As(III) exposure, such as the tyrosine-specific protein phosphatases Ptp2 and Ptp3, modification of which results in their inhibition and consequent activation of Hog1 through the accumulation of basal phosphorylation (Lee and Levin, 2018). Additionally, the glycerol-3-phosphate dehydrogenases, Gpd1 and Gpd2, are inhibited by methylarsenylation in response to As(III) exposure, which prevents glycerol accumulation despite Fps1 closure under this condition (Lee and Levin, 2019). The mechanism that distinguishes a particular cysteine residue as a target of MAs(III) rather than As(III) is not clear. However, in the examples above, the residues modified specifically by MAs(III) are buried within the active sites of the target proteins. It is possible that a hydrophobic environment surrounding the target cysteine residue prevents its modification by As(III), but not by MA(III). Cysteine residues that are targets of As(III) may be solvent exposed and may also be modified by MAs(III).

We provided support for the one-step metabolism model by extending our studies of the MAs(III)-specific target Gpd1. We used a competition assay for in vivo Gpd1-binding by a biotinylated arsenic probe (As-biotin) as an indirect measure of MAs(III) production. We found that pre-treatment of wild-type cells with As(V) did not block As-biotin binding, suggesting that As(V) is not metabolized to physiologically significant levels of MAs(III). However, we were able to force production of MAs(III) from As(V) in an *acr3*Δ mutant, which increases the intracellular As(III) concentration. We were also able to force production of MAs(III) from As(V) by over-expression of the dimeric As(III) methyltransferase Mtq2:Trm112. Thus, it appears that under conditions of As(V) exposure, both free intracellular As(III) and the As(III) methyltransferase are limiting in the production of MAs(III).

### Opposing regulation of Fps1 by arsenylation of Hog1 and methylarsenylation of Acr3

We found additionally that, in response to either As(V) or As(III) exposure, four cysteine residues in the catalytic domain of Hog1 become arsenylated. One effect of Hog1 arsenylation is to prevent its phosphorylation of the Fps1 regulator, Rgc2, and thus to block Fps1 closure by active Hog1. Maintaining Fps1 in an open state when cells are exposed to As(V) appears to confer a growth advantage, as judged by the behavior of a mutant in Hog1 that cannot be arsenylated (*hog1-4C/S*), and which closed Fps1 in response to As(V). Blocking closure of Fps1 through arsenylation of Hog1 appears superficially inconsistent with the finding that As(III) treatment drives Fps1 closure by Hog1. However, we identified a methylarsenylation target that overrides the block imposed by Hog1 arsenylation. We found that Acr3, the plasma membrane As(III) efflux pump, is a component of the Fps1/Rgc1/Rgc2/Hog1 complex and that it additionally serves a novel function as a key regulator of Fps1 channel status in response to arsenic stress. Arsenylated Hog1 is prevented from closing Fps1 by a cytoplasmic loop of Acr3 (loop #4) that appears to interact with Rgc2. However, methylarsenylation of three cysteine residues within this loop in response to As(III) exposure relieves the block to Fps1 closure imposed by Hog1 arsenylation (see Figure 7).

Arsenylation of Hog1 explains its failure to close Fps1 in response to As(V) treatment. However, it is not clear at a mechanistic level how arsenylation of cysteine residues within the Hog1 catalytic domain blocks its ability to phosphorylate Rgc2. The observed effect on Fps1 closure was fully overcome only by mutation to serine of all four arsenylated cysteine residues in Hog1 (data not shown), revealing an aggregate impact of these modifications. It seems unlikely that the effect of Hog1 arsenylation is to prevent its recruitment to Fps1 at the plasma membrane because deletion of the Acr3 cytoplasmic loop #4, which was required for interaction with Rgc2, overcame the block to Fps1 closure by arsenylated Hog1. Indeed, our data suggest that this Acr3 loop is likely to associate with Rgc2 in a manner that shields it from phosphorylation by arsenylated Hog1.

The Acr3 cytoplasmic loop #4 is unique to fungal forms of Acr3-like metalloid transporters (Wawrzycka et al., 2017). We were surprised to find that deletion of this loop conferred greater As(III) tolerance than did wild-type Acr3 (Figure 5), suggesting that Acr3 may be a more effective transporter in the absence of the loop, which may have evolved in fungal species as a functional compromise to block Fps1 closure in response to As(V) challenge.

Our results reveal that stressors with the capability to modify proteins can alter the output of a SAPK. Not only can the SAPK be modified directly in a manner that changes its target specificity, but targets of the SAPK or regulators of those targets may also be modified to alter the SAPK output. Because arsenic has been ubiquitous in the environment since before the emergence of life on earth, organisms have had billions of years to develop defense mechanisms against it. The elegant solution to the problem of distinguishing an As(III) challenge from an As(V) challenge described here reveals an evolutionary adaptation to a toxin whose protein-reactive metabolites are leveraged by the cell for signaling purposes. Although modification of proteins by trivalent arsenicals certainly contributes to their toxicity (Hughes et al., 2011; Martinez et al., 2011), it is also clear that yeast has used this property to craft stress-specific responses to the two major arsenicals in the environment. It seems likely that many other arsenic-modified proteins are integrated into this signaling code.

There is a hint that a version of the signaling code described here may also operate in animals. Methylated metabolites of As(III) (both MAs[III]) and DMA[III]), but not As(III) itself, inhibit the mammalian tyrosine-specific phosphatases PTPB1 and CD45 by reaction with active-site cysteine residues (Rehman et al., 2012). This is similar to our finding that MAs(III) inhibits yeast Ptp2 and Ptp3 to activate Hog1 in response to As(III) treatment (Lee and Levin, 2018). It would be interesting to learn what additional arsenic-binding proteins in human cells react specifically with MAs(III). A previous study of As-biotin binding to proteins in a human proteome microarray revealed an enrichment of glycolytic enzymes (Zhang et al., 2015). These authors suggested that inhibition of glycolysis (and hexokinase-2, in particular) may be key to the anti-cancer properties of As(III) treatment. However, this could be part of an evolved metabolic response to arsenic in mammals that has yet to be elucidated. In any case, this study did not report which of the identified target proteins are bound by As(III) versus MAs(III).

Finally, we have not yet elucidated the pathway by which As(V) activates Hog1, nor the consequences of Hog1 activation in response to As(V) exposure. However, it is clear that this pathway is distinct from that used by As(III), because it stimulates the Hog1 MEK Pbs2 and does not require its metabolism to As(III) (Lee and Levin, 2018). It will be interesting to determine if heretofore unidentified inputs to the HOG pathway are responsible for As(V) activation of Hog1.

## Materials and Methods

### Strains, growth conditions, transformations and gene deletions

The *S. cerevisiae* strains used in this study were all derived from Research Genetics background S288c (Research Genetics, Inc.; Huntsville, AL) and are listed in Table 1. Yeast cell cultures were grown in YPD (1% Bacto yeast extract, 2% Bacto Peptone, 2% glucose) or minimal selective medium, SD (0.67% Yeast nitrogen base, 2% glucose) supplemented with the appropriate nutrients to select for plasmids. Yeast cells were transformed according to Geitz et al. (1995).

**Table 1.**
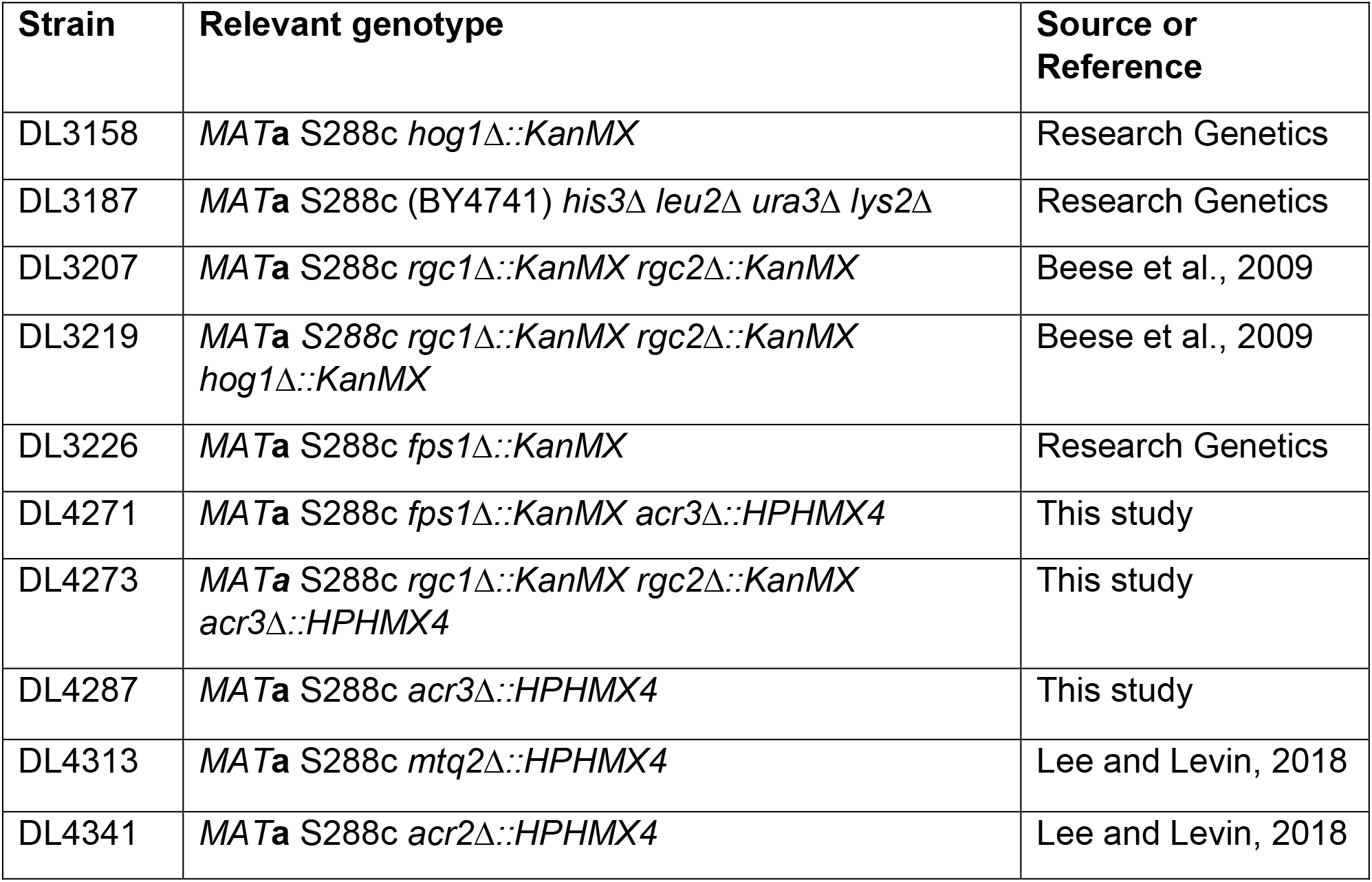
Yeast strains

Chromosomal deletion of the *ACR3* gene was carried out by homologous recombination. The hygromycin-resistance gene *HPHMX4* from pAG32 (Goldstein and McCusker, 1999) was amplified by high-fidelity PCR (Phusion; ThermoFisher; F530S) using primers containing the upstream region (60 bp immediately before the starting ATG) and downstream region (56 bp immediately after the stop codon) of the target gene. The PCR products were integrated into the genome of the wild-type strain by homologous recombination. Integrants were selected on plates containing hygromycin B, yielding *acr3*Δ::*HPHMX4* (DL4287). Deletion of *ACR3* in the *fps1*Δ or *rgc1*Δ*rgc2*Δ background was generated using same method, resulting in *fps1Δ::KanMX acr3Δ::HPHMX4* (DL4271) and *rgc1Δ::KanMX rgc2Δ::KanMX acr3Δ::HPHMX4* (DL4273). All gene replacements were validated by PCR analysis across both integration junctions.

### Chemicals

Sodium arsenite (NaAsO_2_; S7400) and sodium arsenate (Na_2_HAsO_4_; A6756) were purchased from Sigma-Aldrich and methylarsenite (CH_5_AsO_2_; MAs[III]) was purchased from ChemCruz (sc-484527). N-Biotinyl *p*-aminophenyl arsenic acid (As-biotin) was purchased from Toronto Research Chemicals (B394970).

### Plasmid construction and mutagenesis

The *ACR3* gene was epitope-tagged on its C-terminus with the 3xHA epitope and expressed under its native promoter for detection of *ACR3*. The promoter region of *ACR3* (from position -500) and the entire *ACR3* gene without the stop codon was amplified from genomic yeast DNA by high-fidelity PCR (Phusion) using the primers designed with a XhoI site (upstream) and with a NotI site (downstream) and cloned into pRS316-*3HA-ADH1^T^* (p3148) to yield pRS316-*ACR3- 3HA* (p3587).

For co-immunopreciptation (co-IP), Acr3 and Rgc2 were tagged at their C-termini with the 3xHA epitope and expressed under the control of the *MET25* promoter. The *ACR3* coding region was amplified by high-fidelity PCR from genomic yeast DNA using a forward primer that contained an XbaI site (upstream) immediately before the start codon and a reverse primer without a stop codon and a NotI site (downstream) and cloned into pRS316-*MET25*^P^*-RGC2-3HA-ADH1^T^* (p3151) from which Rgc2 was removed by digestion with the same enzymes, to yield pRS316-*MET25*^P^-*ACR3-3HA* (p3470).

For Rgc2 co-IP experiments, the *URA3* marker in pRS316-*MET25*^P^-*RGC2-3HA* was swapped with *HIS3* through homologous recombination. The *pTEF-HIS3-tTEF* sequence from pFA6a-*HIS3MX6* was amplified by high-fidelity PCR using primers containing 40 bp immediately 5’ to the *URA3* promoter in pRS316 and 40 bp immediately 3’ to the *URA3* coding sequence. The PCR products were transformed together with pRS316-*MET25*^P^-*RGC2-3HA* into a wild-type strain, followed by selection on plates for histidine prototrophy. The recombinant plasmid, designated pRS313-*MET25*^P^-*RGC2-3HA* (p3471), was isolated and validated by PCR analysis.

Mtq2 was tagged with the 6xHIS epitope at its C-terminus and expressed under the control of the *MET25* promoter. *MET25^P^-MTQ2-6xHIS* was prepared by double overlap PCR in two steps. First, the *MTQ2* coding sequence was amplified from the genome with a primer that included an overlapping region with *MET25^P^* (upstream) and a primer that included 6xHIS with an XhoI site (downstream). The *MET25^P^* sequence was amplified from pYEp181-*MET25^P^*-*FPS1-Myc* (p3121) with a primer that included a Sac1 site (upstream) and a primer that included an overlapping region with *MTQ2* (downstream). Next, these fragments were amplified together using primers with a Sac1 site (upstream of *MET25^P^*) and an Xho1 site (downstream of *MTQ2-6xHIS*) and cloned into pRS313 vector (p117) at the Sac1 and Xho1 sites to yield pRS313-*MET25^P^-MTQ2- 6xHIS* (p3629).

For Bimolecular Fluorescence Complementation (BiFC) experiments, the *RGC2* coding sequence and the *ACR3* coding sequence were amplified by PCR using primers with BspEI (upstream) and XhoI (downstream) sites for pRS413-CFPC, or XbaI (upstream) and BspEI (downstream) sites for pRS415-CFPN vector, respectively. The digested fragments were cloned into the above vectors, yielding pRS413-*CFPC-RGC2* (p3584) and pRS415-*ACR3-CFPN* (p3585).

Point mutations in Hog1 were generated by QuikChange II mutagenesis (Agilent Technologies; 200523) using template YCplacIII-*HOG1-3HA*. Point mutations and the internal deletion (Δ307-334) in *ACR3* were also generated by QuikChange II mutagenesis using templates, pRS316-*ACR3-3HA,* pRS316-*MET25*^P^-*ACR3-3HA*, or *pRS415-ACR3-CFPN*. Mutant alleles were confirmed by DNA sequence analysis of the entire open reading frame. The plasmids used in this study are listed in Table 2 and the oligonucleotides used are listed in Table 3.

**Table 2.** Plasmids

| **Plasmid** | **Description** | **Source or Reference** |
| --- | --- | --- |
| p115 | pRS316 | Sikorski and Hieter, 1989 |
| p117 | pRS313 | Sikorski and Hieter, 1989 |
| p2492 | pRS426-*MET25<sup>P</sup>-FPS1-Flag* | Beese et al., 2009 |
| p2823 | pAG32 | Goldstein and McCusker, 1999 |
| p3090 | YCplac111-*HOG1-3HA* | F. Posas |
| p3091 | YCplac111 | F. Posas |
| p3121 | pYEp181-*MET25<sup>P</sup>-FPS1-Myc* | Lee et al., 2013 |
| p3148 | pRS316-3HA-*ADH1<sup>T</sup>* | Lee et al., 2013 |
| p3151 | pRS316-*MET25<sup>P</sup>-RGC2-3HA* | Lee et al., 2013 |
| p3155 | pRS316-*MET25<sup>P</sup>-RGC2-S75A,S344A,T808A,S827A,S948A,S1021A,S1035A-3HA (7A)* | Lee et al., 2013 |
| p3182 | pRS316-*RGC2-3HA* | Lee et al., 2013 |
| p3199 | pRS413-*CFPC* | Lipatova et al., 2012 |
| p3200 | pRS415-*CFPN* | Lipatova et al., 2012 |
| p3216 | pRS413-*CFPC-FPS1* (BiFC) | Lee et al., 2013 |
| p3217 | pRS413-*CFPC-HOG1* (BiFC) | Lee et al., 2013 |
| p3225 | pRS426-*HOG1-3HA* | Lee and Levin, 2018 |
| p3460 | pYEP181-*MET25<sup>P</sup>-TRM112* | Lee and Levin, 2018 |
| p3467 | *GAL1<sup>P</sup>-GPD1-TAP* (6HIS-HA-ZZ) | Open Biosystems |
| p3470 | pRS316-*MET25<sup>P</sup>-ACR3-3HA* | This study |
| p3471 | pRS313-*MET25<sup>P</sup>-RGC2-3HA* | This study |
| p3577 | YCplac111-*HOG1-C38S-3HA* | This study |
| p3578 | YCplac111-*HOG1-C156S-3HA* | This study |
| p3579 | YCplac111-*HOG1-C205S-3HA* | This study |
| p3580 | YCplac111-*HOG1-C38S, C156S-3HA* | This study |
| p3581 | YCplac111-*HOG1-C156S, C205S-3HA* | This study |
| p3582 | YCplac111-*HOG1-C38S, C156S, C205S-3HA* | This study |
| p3583 | YCplac111-*HOG1-C38S, C156S, C161S, C205S-3HA* | This study |
| p3584 | pRS413-*CFPC-RGC2* (BiFC) | This study |
| p3585 | pRS415-*ACR3-CFPN* (BiFC) | This study |
| p3586 | pRS415-*ACR3-Δ307-334-CFPN* (BiFC) | This study |
| p3587 | pRS316-*ACR3-3HA* | This study |
| p3588 | pRS316-*ACR3-Δ307-334-3HA* | This study |
| p3589 | pRS316-*ACR3-C316SC, 318S-3HA* | This study |
| p3590 | pRS316-*ACR3-C316S, C318S, C333S-3HA* | This study |
| p3591 | pRS316-*MET25<sup>P</sup> -ACR3-C169S-3HA* | This study |
| p3592 | pRS316-*MET25<sup>P</sup> -ACR3-C333S-3HA* | This study |
| p3593 | pRS316-*MET25<sup>P</sup> -ACR3-C316S, C318S-3HA* | This study |
| p3594 | pRS316-*MET25<sup>P</sup> -ACR3-C316S, C318S, C333S-3HA* | This study |
| p3628 | pRS415-*ACR3-C316S, C318S, C333S-CFPN* (BiFC) | This study |
| p3629 | pRS313-*MET25<sup>P</sup> -MTQ2-6xHIS* | This study |

**Table 3.**
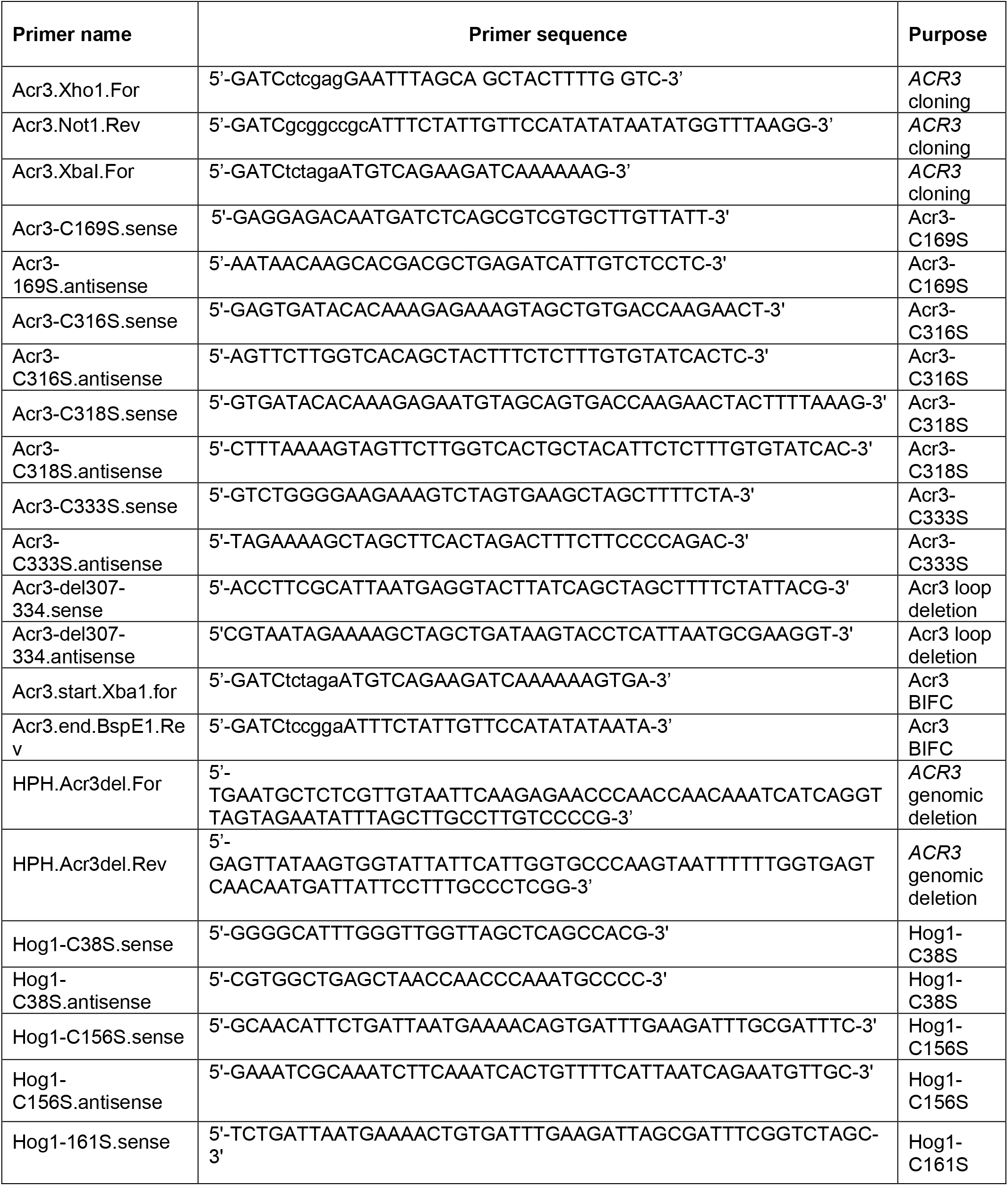
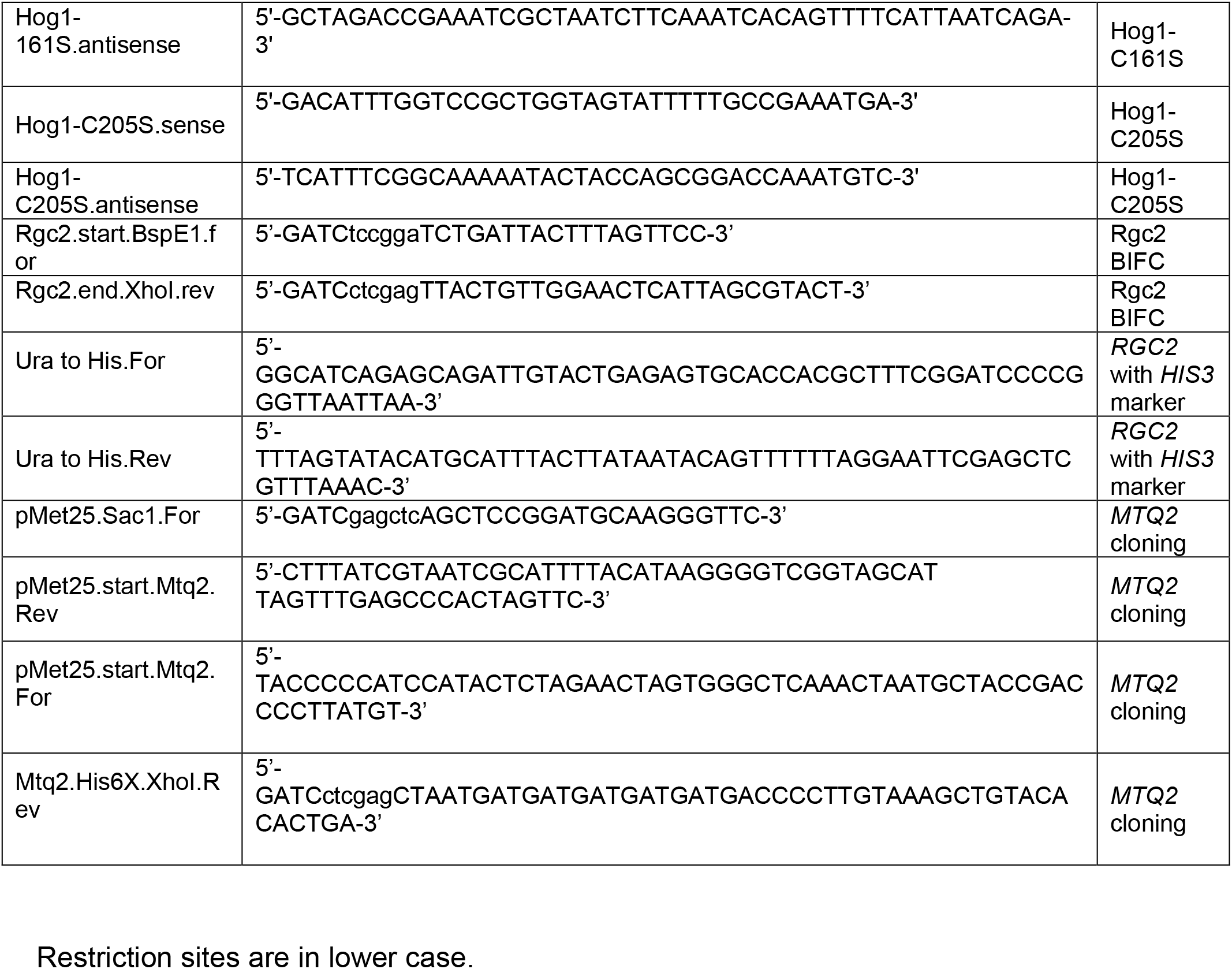
Oligonucleotides

### Protein extraction

Protein extraction for co-immuneprecipitation was carried out as follows. Cells were harvested from 5 ml of medium by centrifugation at 3000g for 5 min. The cell pellet was suspended in 0.7 ml of ice-cold lysis buffer (0.5% Triton X-100, 50 mM Tris-HCl [pH 7.5], 150 mM NaCl, 5 mM EDTA, 5 mM EGTA, 20 µg/ml leupeptin, 20 µg/ml benzamidine, 10 µg/ml pepstatin A, 40 µg/ml aprotinin, 1 mM PMSF, and phosphatase inhibitors [PhosSTOP; Roche; NC0922733]). Glass beads (0.3 mm diam.) were added to this suspension and cells were broken by bead-beating for 1 min at 4 °C. The beads and cell debris were removed by centrifugation at 13,500 rpm for 10 min. at 4 °C and the supernatant was subjected to co-immunoprecipitation. (Kamada et al., 1995; Lee et al., 2013).

Extracts for As-biotin affinity binding experiments were prepared by bead-beating in 50 mM Tris-HCl, pH 7.5, 150 mM NaCl, 0.5% Triton X-100, plus protease inhibitors (cOmplete, Millipore Sigma; 11697498001) (Lee and Levin, 2019).

### Co-immuneprecipitation

Cultures for co-IP experiments (double plasmid transformation) with Fps1-Myc (p3121) and Rgc2-HA (p3151) or Acr3-HA (p3470) were grown to mid-log phase in selective medium and starved for methionine for two hours to induce expression of Fps1, Rgc2 or Acr3, which were expressed under the control of the conditional *MET25* promoter. Cultures for co-IP experiments (triple plasmid transformation) with Fps1-Myc (p3121), Rgc2-HA (p3151) and Acr3 mutants (native promoter) or Fps1-Flag (p2492), Rgc2-HA (p3471) and Hog1 mutants (native promoter) were grown to mid-log phase in selective medium and starved for methionine for three hours to induce expression of Fps1 and Rgc2, which were expressed under the control of the conditional *MET25* promoter. Cultures were then treated with 1mM As(III) or 3mM As(V) for the indicated times.

Co-IPs were carried out as follows: Extracts (100µg of protein) were incubated with mouse monoclonal α-Myc antibody (1µg, 9E10; Pierce; MA1-980), α-HA (Covance; 16B12), or M2 α-Flag antibody (1 µg; Sigma-Aldrich; F3165) for 1 hour at 4°C and precipitated with protein A affinity beads for 1 hour at 4°C. Samples were washed with IP buffer (50 mM Tris-HCl, pH 7.5, 150 mM NaCl, 0.5% Triton) three times and boiled in SDS-PAGE buffer (Lee et al., 2013).

### As-biotin affinity binding experiments

Arsenic-biotin affinity binding experiments were carried out with Gpd1-TAP, Acr3-HA or Hog1-HA. Cultures expressing Gpd1-TAP (under the inducible control of the *GAL1* promoter) were grown in selective medium containing 2% glucose and transferred to selective medium containing 2% raffinose and grown to mid-log phase. Galactose was added to these cultures at a final concentration of 2% to induce expression of Gpd1-TAP for 2 hours. Cultures expressing Acr3-HA (under the repressible control of the *MET25* promoter) were grown to mid-log phase in selective medium and starved for methionine for three hours to induce expression.

Hog1-HA was expressed from the native promoter. Cultures were treated with 10 µM As-biotin for 10 min prior to preparation of extracts. For blocking experiments, cultures were pre-treated with 1 mM As(III), 3 mM As(V) or 0.5 mM MAs(III) for 10 or 20 min and then treated with 10 µM As-biotin for an additional 10 min. Protein extracts (100 µg of protein) were incubated with streptavidin agarose beads (Thermo Scientific) for 1 hour at 4°C. Samples were washed with IP buffer three times and boiled in SDS-PAGE buffer (Lee and Levin, 2019).

For the As-biotin binding assay in wild-type cells over-expressing *MTQ2* and *TRM112*, cultures were triply transformed with *GAL1^P^-GPD1-TAP* (p3467), pYEP181- *MET25^P^-TRM112* (p3460) and pRS313-*MET25^P^-MTQ2-6xHIS* (p3629). Transformants were grown to mid-log phase in selective medium containing 2% raffinose. Cultures were then shifted to methionine-deficient, 2% galactose medium for 2 hours to induce *MTQ2*, *TRM112*, and *GPD1* expression simultaneously.

### SDS-PAGE electrophoresis and immunoblot analysis

Proteins were separated by SDS-PAGE (7.5% or 10% gels) followed by immunoblot analysis on Immobilon-P PVDF membranes (MilliporeSigma; IPVH00005) using mouse monoclonal α-Myc antibody (9E10; Santa Cruz; sc-40), α-HA (16B12; Covance; mms101R), or α-Flag antibody (M2; Sigma-Aldrich; F3165), at a dilution of 1:10,000. Rabbit polyclonal α-phospho-p38 (T180/Y182, Cell Signaling; #9211) was used at a dilution of 1:2,000 to detect phosphorylated Hog1. Secondary goat anti-mouse (Jackson ImmunoResearch; AB_2338447) antibody was used at a dilution of 1:10,000 and secondary donkey anti-rabbit (Amersham; NA9340) was used at a dilution of 1:2,000. Detection of Gpd1-TAP (p1291) was carried out by polyclonal rabbit peroxidase α-peroxidase (PAP; Sigma-Aldrich; P1291) at a dilution of 1:10,000. All results involving immunoblot analyses were replicated at least once and representative blots are shown.

### Bimolecular Florescence Complementation (BiFC)

Wild-type or mutant haploid cells were transformed with combinations of plasmids that express Acr3-CFPN (pRS415*-ACR3*-*CFPN*; p3585), or its mutant form (pRS415*-ACR3*-*Δ307-334-CFPN*; p3586), CFPC-Fps1 (pRS413-*CFPC-FPS1*; p3216), CFPC-Rgc2 (pRS413-*CFPC-RGC2*; p3584), CFPC-Hog1 (pRS413-*CFPC-HOG1*; p3217), or empty vectors (pRS413-*CFPC* or pRS415-CFPN; p3199 or p3200). Transformants were grown overnight in SD medium, then diluted in YPD for growth to mid-log phase, centrifuged and resuspended in SD medium. Cells were visualized at 23°C with a Zeiss Axio Observer Z1 with a Zeiss Plan-Apochromat 100x/1.4 oil-immersion objective, fitted with a CFP filter, or by differential interference contrast (DIC), and photographed with a Hamamatsu Orca R2 CCD Camera. Images were processed using Zen Pro software.

### Measurement of intracellular glycerol concentrations

Intracellular glycerol concentrations were measured in whole cells grown in YPD, treated with 3 mM As(V) for two hours, and centrifuged briefly to remove the culture supernatant. Enzymatic assays for glycerol were carried out using a kit from R-Biopharm (10148270035) and normalized to A_600_ of the initial culture. The mean and standard deviation from three independently grown cultures was presented for each value.

### Growth curve assay

Colonies from a *hog1*Δ strain (DL3158), transformed with centromeric plasmids bearing *HOG1* (p3090), *hog1-4C/S* (p3583), or vector (p3091), were grown in selective medium to mid-logarithmic phase and treated with 1 mM As(V) and growth was followed by spectrophotometry. All values for doubling times represent the mean from three independent transformants.

## Notes on reproducibility

All immunoblots, co-IPs, and As-biotin pull-downs were reproduced at least once in independent experiments with representative images shown.

## Supplemental material

**Figure S1.** The *hog1-4C/S* mutant induces dissociation of Rgc2 from Fps1 normally in response to As(III) exposure.

**Figure S2.** Methylarsenite pre-treatment diminishes As-biotin binding to Hog1.

**Figure S3.** The *hog1-4C/S* mutant does not display hyper-sensitivity to As(V).

**Figure S4. T**he association between Acr3 and Fps1 is unaffected by exposure to either As(III) or As(V).

**Figure S5.** Bimolecular Fluorescence Complementation controls.

## Acknowledgments

This work was supported by a grants from the NIH (R01GM48533 and R01GM138413) to DEL.

## Author Contributions

D. E. Levin and J. Lee contributed to the design of the experimental approach, interpretation of data, and writing the manuscript. J. Lee conducted all of the experiments.

## Supplemental Figure legends

**Supplemental Figure S1.**
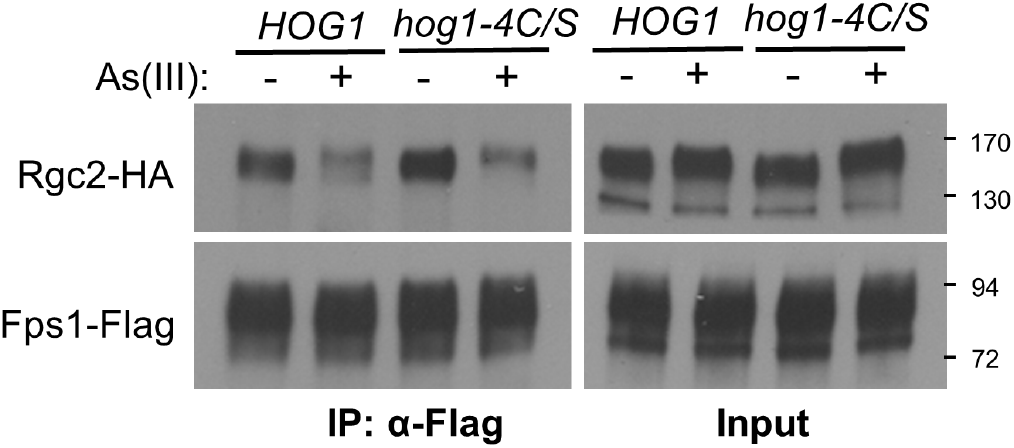
The *hog1-4C/S* mutant induces dissociation of Rgc2 from Fps1 normally in response to As(III) exposure. A *hog1*Δ strain (DL3158) co-transformed with plasmids expressing Rgc2-HA (p3471), Fps1-Flag (p2492), and either wild-type Hog1 (p3090) or Hog1-4C/S (p3583) was either treated with 1 mM As(III) for 10 min. or not (-) prior to processing for co-IP with anti-Flag antibodies.

**Supplemental Figure S2.**
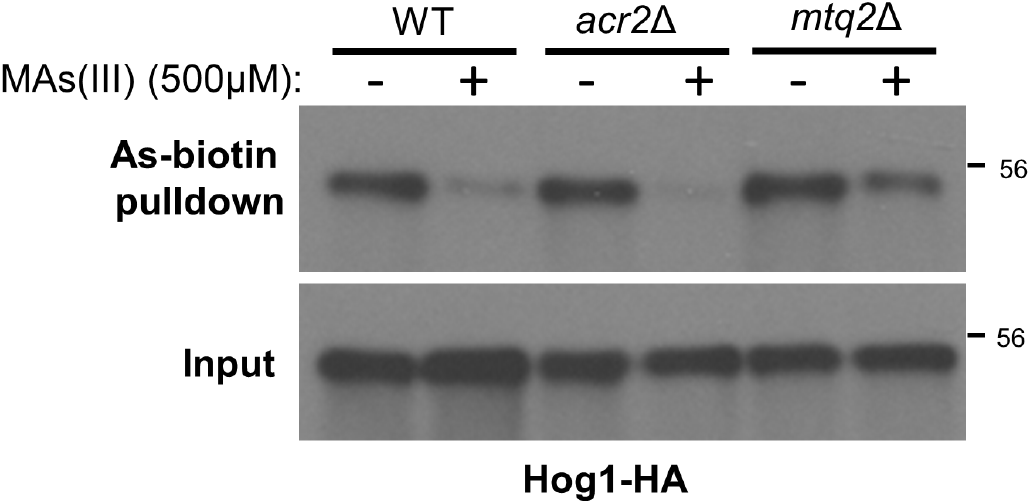
Methylarsenite pre-treatment diminishes As-biotin binding to Hog1. Wild-type strain (DL3187), *acr2*Δ (DL4341), or *mtq2*Δ (DL4313) strains were transformed with a multi-copy plasmid expressing Hog1-HA (p3225) and treated (+) or not (-) with 500 μM MAs(III) for 20 min prior to treatment with 10 μM As-biotin for 10 min. Extracts were subjected to affinity pull-down with SA beads prior to SDS-PAGE and immunoblot analysis for Hog1-HA.

**Supplemental Figure S3.**
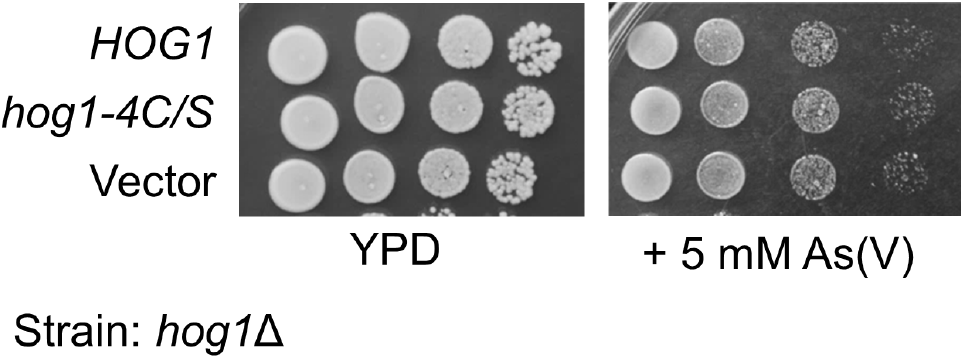
The *hog1-4C/S* mutant does not display hyper-sensitivity to As(V). Cultures of a *hog1*Δ strain (DL3158) transformed with a centromeric plasmid expressing the *hog1-4C/S* allele (p3583), or vector (p3090), were spotted onto YPD, YPD plus 5mM As(V), at serial 10-fold dilutions (from left to right) and incubated at 30°C for 3 days.

**Supplemental Figure S4.**
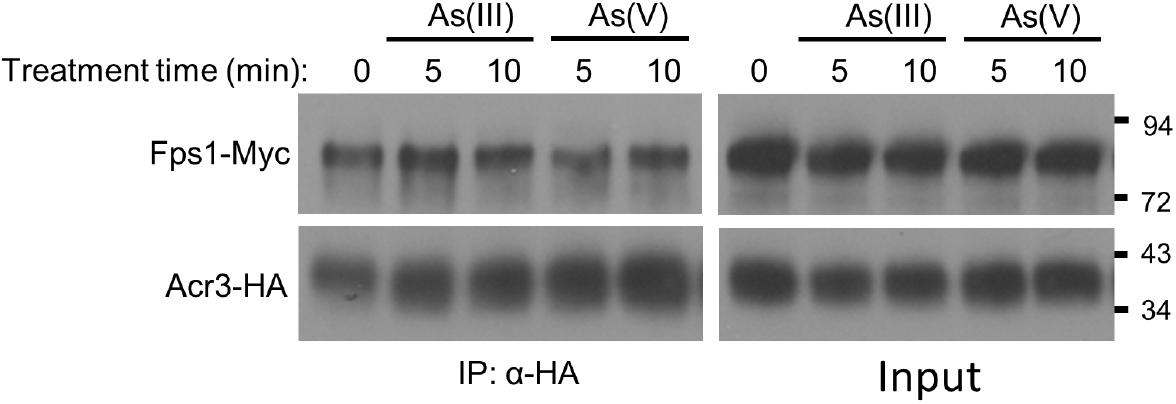
The association between Acr3 and Fps1 is unaffected by exposure to either As(III)or As(V). Rgc2-HA and Fps1-Myc were co-expressed (from p3151 and p3121, respectively) in a wild-type strain (DL3187). The strain was treated with 1 mM As(III) or 3 mM As(V) for the indicated times and processed for co-IP of Fps1 with Acr3 using anti-HA antibodies. Molecular mass markers (in kDa) are shown on the right.

**Supplemental Figure S5.**
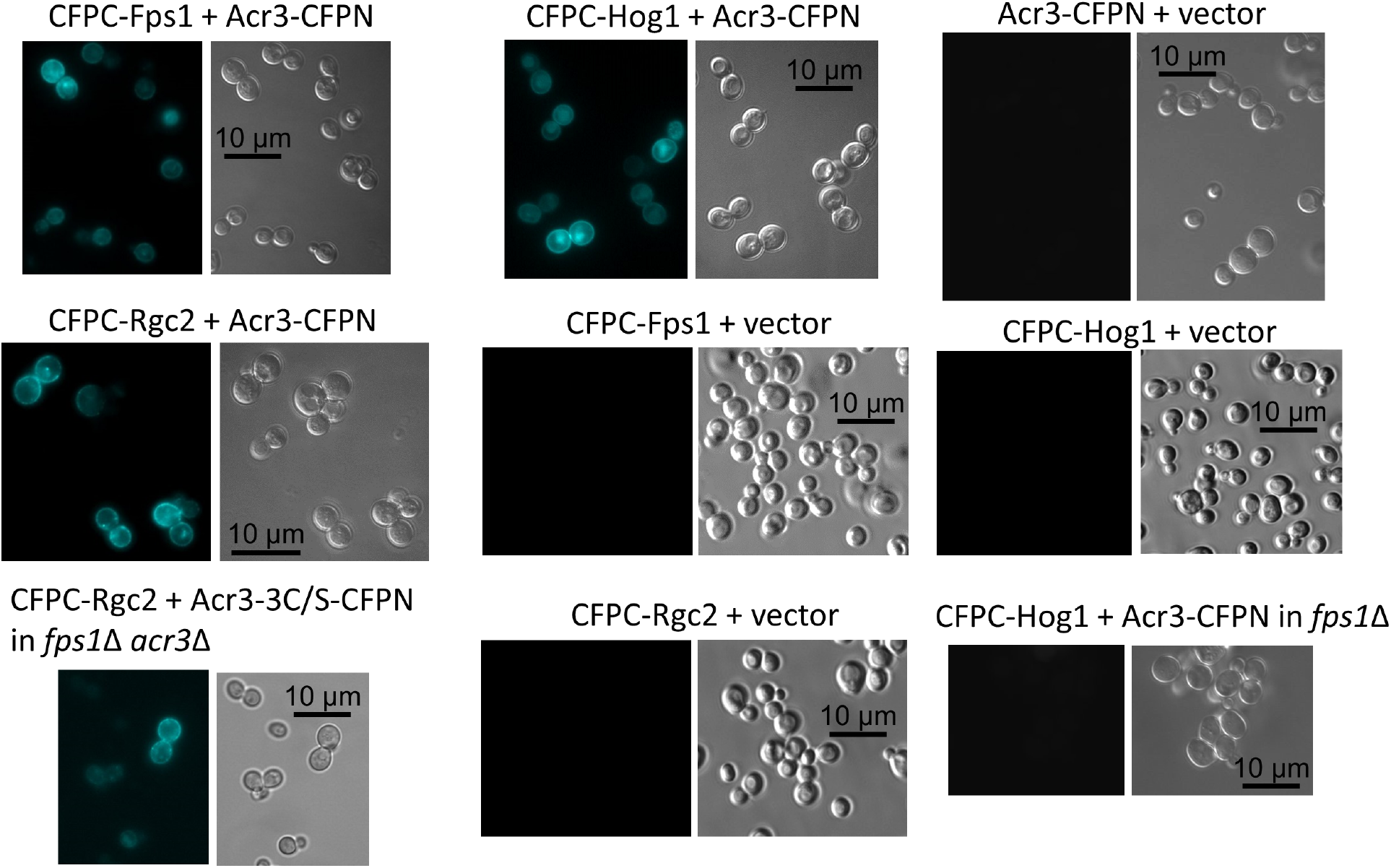
Bimolecular Fluorescence Complementation controls. A wild-type strain (DL3187), an *fps1*Δ mutant (DL3226), or an *acr3*Δ *fps1*Δ mutant (DL4271) were transformed with the indicated BiFC plasmids and visualized under a UV light source to reveal fluorescence complementation (left), or under visible light (DIC; right). Plasmids expressed: CFPC-Fps1 (p3216), CFPC-Hog1 (p3217), CFPC-Rgc2 (p3584), Acr3-CFPCN (p3585), Acr3-3C/S-CFPN (p3628), or vectors (p3199 or p3200).

